## Supplementary material for "Topological Environment in Genetic and Metabolic Networks": Support information

**Allosteric system for length of the pathway $n = 3$ in End-Product Inhibition.**

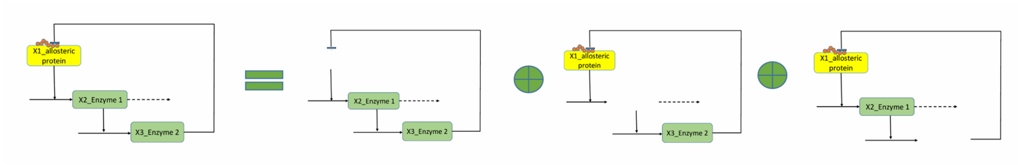


Figure 1: The allosteric system of end-product inhibition for a pathway of length $n = 3$ serves as a model for biosynthetic pathways (see [5]), such as the biosynthesis of homoserine regulated by methionine (see EcoCyc pathways platform). The topological environment of this system is divided into three local charts, corresponding to three Hilbert bases.

**The Power-Law Model is governed by (see [5]):**

$$\frac{dX_1}{dt} = D^{-1}(\alpha_{Max} + \alpha_{Min} X_3^g K_R^{-g}) - \beta_1 X_1 \tag{1}$$

$$\frac{dX_2}{dt} = \alpha_2 X_1 - \beta_2 X_2 \tag{2}$$

$$\frac{dX_3}{dt} = \alpha_3 X_2 - \beta_3 X_3 \tag{3}$$

Recasting:

$$D = 1 + K_R^{-g} X_3^g \tag{4}$$

We define the support spaces for each one of the equations:

$$Supp(\dot{X}_1) = \{(1,0,0,0), (0,0,g,-1), (0,0,0,-1)\}, \tag{5}$$

$$Supp(\dot{X}_2) = \{(1,0,0,0), (0,1,0,0)\}, \tag{6}$$

$$Supp(\dot{X}_3) = \{(0,1,0,0), (0,0,1,0)\}. \tag{7}$$

We note that the lattice vectors are ordered in lexicographic order. That is, in $Supp(\dot{X}_1)$, the vector $(1,0,0,0)$ satisfies $(1,0,0,0) \geq_{lex} (0,0,g,-1)$ if and only if the subtraction of these vectors has, in its first nonzero coordinate, a value greater than or equal to zero. For example,

$$(1,0,0,0) - (0,0,g,-1) = (1,0,-g,1),$$

whose first coordinate is 1, which is positive. Therefore, these vectors are ordered lexicographically. Thus, the following order holds in $Supp(\dot{X}_1)$:

$$(1,0,0,0) \geq_{lex} (0,0,g,-1) \geq_{lex} (0,0,0,-1).$$

Similarly, the sets $Supp(\dot{X}_2)$ and $Supp(\dot{X}_3)$ are also ordered lexicographically.

**Topologival Environment and Orthogonal Enzyme Matrices.**

In the following, we compute the lattice cones from the exponent space (*supp*) for each differential equation. We also construct the corresponding orthogonal matrices. The **Topological Environment** is encoded in this set of orthogonal matrices. In simple terms, a numerical orthogonal matrix is associated with each one. The values are determined according to the neighboring nodes or neighboring enzyme reactions, as shown below.

$$Con(\dot{X}_1) = \begin{pmatrix} 1 & 0 & 0 & 0 \\ 1 & 0 & -g & 0 \\ 0 & 0 & g & 0 \\ 0 & 0 & 0 & -1 \end{pmatrix}$$

In the topological environment for $\dot{X}_1$, we observe that the neighboring enzymes in the first row $(1,0,0,0)$ of the cascade network are the enzymes $X_2$ and $X_3$. Thus, an orthogonal vector in the corresponding first row of the orthogonal matrix could be $(0,1,1,0)$. For the second row in $Con(\dot{X}_1)$, i.e., $(1,0,-g,0)$, the only neighboring enzyme to the proteins $X_1$ and $X_3$ is $X_2$. Therefore, an orthogonal vector could be $(0,1,0,0)$. We selected the orthogonal vectors in the simplest way. This is justified because the Hilbert bases are the minimal generators of these lattice vectors. Thus, the last two vectors in $Con(\dot{X}_1)$ were chosen.

The central core of this methodology is based in the orthogonal matrices, **they codify a local dynamics for each local chart through the Hilbert basis**.

**Toplogical environment coded for the allosteric enzyme $X_1$.**

$$Con^{\vee}(\dot{X}_1) = \begin{pmatrix} 0 & 1 & 1 & 0 \\ 0 & 1 & 0 & 0 \\ 1 & 1 & 0 & 0 \\ 1 & 0 & 0 & 0 \end{pmatrix}$$

$$Con(\dot{X}_2) = \begin{pmatrix} 1 & 0 & 0 & 0 \\ 1 & -1 & 0 & 0 \\ 0 & 1 & 0 & 0 \end{pmatrix}$$

**Toplogical environment coded for the allosteric enzyme $X_2$.**

$$Con^{\vee}(\dot{X}_2) = \begin{pmatrix} 0 & 1 & 0 & 0 \\ 0 & 0 & 1 & 0 \\ 1 & 0 & 1 & 0 \end{pmatrix}$$

$$Con(\dot{X}_3) = \begin{pmatrix} 0 & 1 & 0 & 0 \\ 0 & 1 & -1 & 0 \\ 0 & 0 & 1 & 0 \end{pmatrix}$$

**Toplogical environment coded for the allosteric enzyme $X_3$.**

$$Con^{\vee}(\dot{X}_3) = \begin{pmatrix} 1 & 0 & 1 & 0 \\ 1 & 0 & 0 & 0 \\ 1 & 1 & 0 & 0 \end{pmatrix}$$

$$Con(D) = \begin{pmatrix} 0 & 0 & g & 0 \end{pmatrix}$$

It is possible that topological environmnet for $D$ be the same for $X_3$. Because, this variable is in function with it.

**Hilbert bases are associated with each lattice cone in the Topological Environment. For the computation, one can use platforms in Computational Algebraic Geometry such as `SageMath` or `Normaliz`. For the cases of End-Product Inhibition, the following tutorial has been developed in [6], [3].** The Hilbert basis for this system can be the identity matrix or permuted identity matrices, such that the Hilbert basis are invariant under linear transformations, such as permutations on the rows.

$$\begin{pmatrix} 1 & 0 & 0 & 0 \\ 0 & 1 & 0 & 0 \\ 0 & 0 & 1 & 0 \\ 0 & 0 & 0 & 1 \end{pmatrix} \text{Permuting rows, 1 with 2} \begin{pmatrix} 0 & 1 & 0 & 0 \\ 1 & 0 & 0 & 0 \\ 0 & 0 & 1 & 0 \\ 0 & 0 & 0 & 1 \end{pmatrix}$$

$$\begin{pmatrix}1&0&0&0\\0&1&0&0\\0&0&1&0\\0&0&0&1\end{pmatrix} \text{ Permuting rows, 1-2 and 2-3 } \begin{pmatrix}0&1&0&0\\0&0&1&0\\1&0&0&0\\0&0&0&1\end{pmatrix}$$

$$\begin{pmatrix}1&0&0&0\\0&1&0&0\\0&0&1&0\\0&0&0&1\end{pmatrix} \text{ Permuting rows, 1-2, 2-3, 3-4 } \begin{pmatrix}0&1&0&0\\0&0&1&0\\0&0&0&1\\1&0&0&0\end{pmatrix}$$

$$\begin{pmatrix}1&0&0&0\\0&1&0&0\\0&0&1&0\\0&0&0&1\end{pmatrix} \text{ Permuting rows, 1 with 2 } \begin{pmatrix}1&0&0&0\\0&0&1&0\\0&1&0&0\\0&0&0&1\end{pmatrix}$$

Also, it is possible to predict values for the number of binding sites using a topological environment. We consider **the total sum of these permuted matrices, this matrix could be a gene matrix interaction:**

$$\text{M(Total)=}\begin{pmatrix}2&2&0&0*\\1&0&3&0\\1&1&1&1*\\1&0&0&3*\end{pmatrix}$$

We multiply the informative lattice vector $(1, 0, -g, 1)$ row by row, only by the rows in asterisk, and hence.

$$(2, 2, 0, 0) * (1, 0, -g, 1) = 2 \geq 0,$$

$$(1, 1, 1, 1) * (1, 0, -g, 1) = 1 - g + 1 \geq 0,$$

$$(1, 0, 0, 3) * (1, 0, -g, 1) = 1 + 3 \geq 0,$$

Add all the inequalities. It yields: $8 - g \geq 0$, $\mathbf{g \leq 8}$. Therefore, we take $\mathbf{g = 8}$, as is shown in [5].

We have chosen only the Hilbert basis for $X_1$. That is why, this is the only non linear expression. The expressions for

$$and$$

are linear. Thus **Topological evironment only is useful for $X_1$.**

$$HB1 = \begin{pmatrix} 0 & 1 & 0 & 0 \\ 1 & 0 & 0 & 0 \\ 0 & 0 & 1 & 0 \\ 0 & 0 & 0 & 1 \end{pmatrix}$$

The monomial transformation, regard with this methodology, is given by: $X_1 = u_2$, $X_2 = u_1$, $X_3 = u_3$, $X_4 = u_4$.

**Fix Points on Torus are :**

At the same time, We only use the formula and the elements of Hilbert basis for fix points on torus for $X_1$.

Here, the symbols for Hilbert basis are: $h_1, h_2, h_3, h_4$, and the rows in the lattice-cone $Con(\dot{X}_1)$ are: $b_1, b_2, b_3, b_4$. Thus the fix point on torus are computed, as follows:

$$X_{b_1} = \lim_{Z \longrightarrow 0} Z^{<b_1,h_1>} = \lim_{Z \longrightarrow 0} (Z^{<b_1,h_1>}, Z^{<b_1,h_2>}, Z^{<b_1,h_3>}, Z^{<b_1,h_4>}). \quad (8)$$

$$X_{b_1} = \lim_{Z \longrightarrow 0} (Z^{<(1,0,0,0),(0,1,0,0)>}, Z^{<(1,0,0,0),(1,0,0,0)>}, Z^{<(1,0,0,0),(0,0,1,0)>},$$

$$Z^{<(1,0,0,0),(0,0,0,1)>}).(9)$$

$$X_{b_1} = \lim_{Z \longrightarrow 0} (Z^0, Z^1, Z^0, Z^0) = \lim_{Z \longrightarrow 0} (1, Z^1, 1, 1) = (1, 0, 1, 1). \quad (10)$$

$$X_{b_2} = \lim_{Z \longrightarrow 0} (Z^{<(1,0,-g,1),(0,1,0,0)>}, Z^{<(1,0,-g,1),(1,0,0,0)>}, Z^{<(1,0,-g,1),(0,0,1,0)>},$$

$$Z^{<(1,0,-g,1),(0,0,0,1)>}).(11)$$

$$X_{b_2} = \lim_{Z \longrightarrow 0} (Z^0, Z^1, Z^{-g}, Z^1) = (1, 0, \infty, 0). \quad (12)$$

$$X_{b_3} = \lim_{Z \longrightarrow 0} (Z^{<(0,0,g,0),(0,1,0,0)>}, Z^{<(0,0,g,0),(1,0,0,0)>}, Z^{<(0,0,g,0),(0,0,1,0)>},$$

$$Z^{<(0,0,g,0),(0,0,0,1)>}).(13)$$

$$X_{b_3} = \lim_{Z \longrightarrow 0} (Z^0, Z^0, Z^g, Z^0) = (1, 1, 0, 1). \quad (14)$$

$$X_{b_4} = \lim_{Z \longrightarrow 0} (Z^{<(0,0,0,0,-1),(0,1,0,0)>}, Z^{<(0,0,0,0,-1),(1,0,0,0)>}, Z^{<(0,0,0,0,-1),(0,0,1,0)>},$$

$$Z^{<(0,0,0,0,-1),(0,0,0,1)>}).(15)$$

$$X_{b_4} = \lim_{Z \longrightarrow 0} (Z^0, Z^0, Z^0, Z^{-1}) = (1, 1, 1, \infty). \quad (16)$$

We will choose the fixed point on the torus $(u_1, u_2, u_3, u_4) = (1, 1, 0, 1)$ to linearize our system of equations and study the local dynamics. The system of differential equations in the new system of monomial coordinates is given by:

$$\dot{X}_2 = \dot{u}_1 = \alpha_2 u_2 - \beta_2 u_1; \quad (17)$$

$$\dot{X}_1 = \dot{u}_2 = -\beta_1 u_2 + g\alpha_{MIN} K^{-g} u_3 - u_4(\alpha_{Max} + \alpha_{Min} K^{-g}); \quad (18)$$

$$\dot{X}_3 = \dot{u}_3 = \alpha_3 u_1 - \beta_3 u_3; \quad (19)$$

Firstly. it will be useful to linearize the equation for the constraint $D$ through the Jacobian matrix, as follows:

$$J(\frac{D(u1, u2, u3)}{(u1, u2, u3)})_{(1,1,0,1)} * U = \begin{pmatrix} 0 & gK_R^{-g} u_2^{g-1} & 0 & 0 \end{pmatrix} * U = gK_R^{-g} u_2.$$

In addition, we also proceed to linearize the system of differential equations using the new system of monomial coordinates $U = (u_1, u_2, u_3)$. Substituting the linearized equation for $D$, we obtain:

$$\dot{u}_1 = \alpha_2 u_2 - \beta_2 u_1; \quad (20)$$

$$\dot{u}_2 = -\beta_1 u_2 + g\alpha_{MIN} K^{-g} u_3 - gK_R^{-g} u_2 * (\alpha_{Max} + \alpha_{Min} K^{-g}); \quad (21)$$

$$\dot{u}_3 = \alpha_3 u_1 - \beta_3 u_3; \quad (22)$$

Hence, is obtained a classical linear system of diferential equations, such as:

$$\dot{U} = A * U^T; \quad (23)$$

With parametres defined as:

$$A = \begin{pmatrix} -A & B & 0 \\ 0 & -C & D \\ E & 0 & -F \end{pmatrix}$$

$A = \beta_2$, $B = \alpha_1$, $C = -\beta_1 - gK_R^{-g}\,(\alpha_{\text{Max}} + \alpha_{\text{Min}} K^{-g})$, $D = g\,\alpha_{\text{Min}} K^{-g}$, $E = \alpha_3$, $F = \beta_3$.

The classical analysis of this allosteric system with end-product inhibition is presented in [**?**].
The condition for sustained oscillatory stability in this system is given by:

$$ACF >> BDE \tag{24}$$

### 1 Differences in Classical fixed points in dynamical systems.

Classical fixed points: Any solution $X* \in \mathbb{C}^n$ or $\mathbb{R}^n$ where $f(x*)$, including zeros.

Torus fixed points: Only those equilibria where each coordinate is nonzero (since the torus excludes zero).

So, some fixed points from the classical sense might disappear in the torus setting if they involve zeros.

Conversely, on the torus, you often express fixed points in terms of monomial/logarithmic coordinates, which can reveal structure (e.g., oscillations, scaling laws) that is not obvious in the additive classical coordinates.

**We remark in this point, that fix points on torus can reveal new oscillatory behaviors in Biology, modeled through Power-law for genetic and metabolic Networks.**

To continuation to analyse more this conjeture, we give a formal proof, that sustain deep relationship between formal polynomial like Power-Law, their lattice-cones and toric varieties associated. With this fact, we assert that the classical fix points modeled by S-systems equations are contained in a class more general of objects, those are, fix points on algebraic torus, A fix point here, corresponds to a toric ideals, or a binomaial generator, to this generator belongs the S-systems equations.

**A Proof for the commutativity of the diagram.**

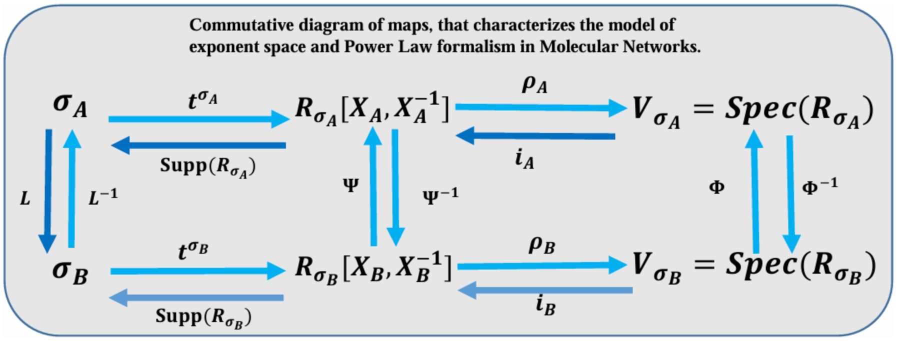


Figure 2: Commutative diagram of mappings between the exponent space, the ring of power-law polynomials for molecular networks, and the prediction of the number of binding sites. In formal mathematics, a diagram of maps is commutative if the composition of maps is consistent across all pathways. Each square in the diagram must commute, as explicitly we can see: $t^{\sigma_A}(L^{-1}) = \Psi(t^{\sigma_B})$, and, $t^{\sigma_B}(L) = \Psi^{-1}(t^{\sigma_A})$. Similarly, the second square must also commute: $i_A(\Phi) = \Psi(i_B)$, and $i_B(\Phi^{-1}) = \Psi^{-1}(i_A)$. The definitions for, $t^{\sigma_A}$, $L$, $L^{-1}$, $\Psi$, $\Phi$, $i_A$, $i_B$, $R_{\sigma_A}$, and $Spec(R_{\sigma_A})$, $Supp(R_{\sigma_A})$ are provided in the Methods section. We assert that this diagram is commutative; see complete proof in, [2, 4] and supporting information.

**A sketch of the proof.** Let $h_{\sigma_A} : \sigma_A \longrightarrow R_{\sigma_A}$, $h_{\sigma_B} : \sigma_B \longrightarrow R_{\sigma_B}$, $\rho_A : R_{\sigma_A} \longrightarrow V_{\sigma_A}$ and $\rho_B : R_{\sigma_B} \longrightarrow V_{\sigma_B}$ be monomial homomorphisms. By hypothesis, the lattice cones $\sigma_A$ and $\sigma_B$ are isomorphic, that is, $\sigma_A \approx \sigma_B$. Then there exists an unimodular linear transformation given by, $L$, such that $L(\sigma_A) = \sigma_B$ and vice versa $L^{-1}(\sigma_B) = \sigma_A$. This linear transformation is well defined through the Hilbert basis. With this fact, we build the monomial homomorphisms, as follows,

$$h_{\sigma_A}(a) = \sum \lambda_a t^a \in R_{\sigma_A} \text{ and } a \in supp(h_{\sigma_A}) \subset \sigma_A,$$

$$h_{\sigma_B}(b) = \sum \lambda_b t^b \in R_{\sigma_B} \text{ and } b \in supp(h_{\sigma_B}) \subset \sigma_B,$$

$\Psi(h_{\sigma_B}) = \sum \lambda_B t^{L(\sigma_A)} \in R_{\sigma_A}$ and $L(\sigma_A) = \sigma_B \in supp(\Psi^{-1}) \subset \sigma_B$,

$\Psi^{-1}(h_{\sigma_A}) = \sum \lambda_A t^{L^{-1}(\sigma_B)} \in R_{\sigma_B}$ and $L^{-1}(\sigma_B) = \sigma_A \in supp(\Psi) \subset \sigma_A$.

We also chose the prime generators $t^a \in R_{\sigma_A}$, $a \in \sigma_A$, $t^b \in R_{\sigma_B}$, and $b \in \sigma_B$. and We define the following maps:

$\rho_A(t^a) = < t^a > \in V_{\sigma_A} = Spec(R_{\sigma_A})$,

$\rho_B(t^b) = < t^b > \in V_{\sigma_B} = Spec(R_{\sigma_B})$,

$\Phi(< t^b >) = < t^{L^{-1}(b)} > \in V_{\sigma_A} = Spec(R_{\sigma_A})$,

$\Phi^{-1}(< t^a >) = < t^{L(a)} > \in V_{\sigma_B} = Spec(R_{\sigma_B})$.

Here, the $< t^a >$ and $< t^{L^{-1}(b)} >$ are prime ideals, from the spectrum of coordinate rings, of Laurent´s formal power series $R_{\sigma_A}$ and $R_{\sigma_B}$ respectively. We can see without lost of generality that the monomial homomorphisms fulfill the following identities, we can take, $\lambda_a = \lambda_b = 1$, hence, $L \circ L^{-1} = id_{\sigma_B}$, $L^{-1} \circ L = id_{\sigma_A}$, $\Psi \circ \Psi^{-1} = id_{R_{\sigma_A}}$, $\Psi^{-1} \circ \Psi = id_{R_{\sigma_B}}$, and $\Phi \circ \Phi^{-1} = id_{V_{\sigma_A}}$, $\Phi^{-1} \circ \Phi = id_{V_{\sigma_B}}$. We remember that the maps, $L$, $\Psi$, and $\Phi$ are isomorphisms for, lattice cones, algebras of coordinate rings, and toric morphisms. The two first morphisms are well defined, one is an unimodular transformation, second one is an algebra homomorphism. Only we need to show that $\Phi$ is a toric morphism. We define the monomial homomorphism $\Phi^* : \mathbb{C}^n \longrightarrow \mathbb{C}^n$ such that, $\Phi^*(< t^b >) =< t^a > \ni \Phi^*(V_{\sigma_B}) \subset V_{\sigma_A}$ this homomorphism induces the morphism $\Phi$ it which is bijective, since that for the generator, $t^0 = id_{V_{\sigma_B}}$, with lattice vector $a = 0 \in \sigma_A$, $\Phi(t^0) = t^{L^{-1}(0)} = id_{V_{\sigma_A}}$. Therefore $\Phi$ is injective, Now we take the generator $t^b \in V_{\sigma_B}$, thus $L(a) = b$, $\Longrightarrow \exists\, t^a \in V_{\sigma_A} \ni \Phi(t^b) = t^{L^{-1}(b)} = t^a$, therefore $\Phi$ is sobrejective. We can notice that we can write, $\Phi = \Phi^*|_{V_{\sigma_B}}$. At the same time, we can notice that $\Phi^{-1}$ is also a toric morphism. For the map $\Psi$, we can see easily that, $\Psi(h_{\sigma_B}(b)) = h_{\sigma_A}(L^{-1}(b))$ and $\Psi(i_B(< t^b >)) = i_A(< t^{L^{-1}(b)} >)$. The proof for the isomorphisms $V_{\sigma_B} \approx V_{\sigma_A}$ implies the isomorphism $\sigma_B \approx \sigma_A$, see Ewald ¡[4]. Together, all those facts proved the commutativity of the diagram in Figure 2, **q.e.d.**

**A proof for the theorem 3.3 in the manuscript.**

**Theorem: (Toric S-system)**: Let

$$\dot{X}_i = \alpha_{ip_i} \prod_{j=1}^{m} X_j^{g_{ip_i}} - \beta_{iq_i} \prod_{j=1}^{m} X_j^{h_{iq_i}}, \quad i = 1, \ldots, n,$$

be a set of S-system equations. Then, there exists a toric ideal, namely

$$I_H = \langle X^a - X^b \ : \ H(a-b)^T = 0 \in \mathbb{Z}^n, , a, b \in \mathbb{Z}^n \ \rangle,$$

where $H$ is a Hilbert basis with $\det(H) = \pm 1$. Moreover, the toric binomials $(X^a - X^b)$ generate the S-system equations, that is, $\dot{X}_i \in I_H$.

**Proof:** We construct the exponent space or lattice cone from the support sets for the polynomials $\dot{X}_i$, for some equation of the $i$ S-system, i.e., $Con(\dot{X}_i) = (g_{ip_i}, .., g_{ip_i} - h_{iq_i}, ..., h_{iq_i})$, where the lattice vectors $(g_{ip_i} - h_{iq_i})$ are the pairwise differences in lexicographic order, with the propose to build a convex geometry, thus we apply the algorithm of Hilbert basis over this lattice cone to find basis, we write in symbols this fact,
$H = \{h_1, ..., h_k\}$, we give a matrix representation of it if we consider each $h_i$ as a row vector; thus we write $h_1 = (u_{11}, u_{12}, ..., u_{1n})$, $h_2 = (u_{21}, u_{22}, ..., u_{2n})$,...,$h_k = (u_{k1}, u_{k2}, ..., u_{kn})$.

$$H = \begin{pmatrix} u_{11} & u_{12} & ... & u_{1n} \\ u_{21} & u_{22} & ... & u_{2n} \\ ... & ... & ... & ... \\ u_{k1} & u_{k2} & ... & u_{kn} \end{pmatrix}$$

To continuation, we can use the matrix defined above to represent the variables $X_i$ in a new system of coordinates, a using the monomial transformation, $u_{ij} \longmapsto w_i^{u_{ij}}$, see [4], the relevance to do that, is to notice dynamics properties encrypted for S-systems into of the Hilbert basis.

$$\begin{array}{rlll} X_1 = w_1^{u_{11}} * & w_2^{u_{12}} * & ... & * w_n^{u_{1n}}, \\ X_2 = w_1^{u_{21}} * & w_2^{u_{21}} * & ... & * w_n^{u_{2n}} \\ ... & ... & ... & ... \\ X_k = w_1^{u_{k1}} * & w_2^{u_{k2}} * & ... & * w_n^{u_{kn}} \end{array} \tag{25}$$

If we substitute this change of coordinates in the original S-systems, we have the following equation, it which is valid for all $i = 1, .., n$,

$$\dot{X}_i(w_1, ..., w_k) = \alpha_{ip_i} \prod_{j=1}^{m} (w_j^{u_{j1}} * w_2^{u_{j2}} * ... * w_n^{u_{jn}})^{g_{ip_i}} - \beta_{iq_i} \prod_{j=1}^{m} (w_j^{u_{j1}} * w_2^{u_{j2}} * ... * w_n^{u_{jn}})^{h_{iq_i}}. \tag{26}$$

Or simply,

$$\dot{X}_i(w_1, ..., w_k) = \alpha_{ip_i} \prod_{j=1}^{m} (W_i^{h_j})^{g_{ip_i}} - \beta_{iq_i} \prod_{j=1}^{m} (W_i^{h_j})^{h_{iq_i}}. \tag{27}$$

Now, we build the new lattice vector for this $i$-esim S-system is: $(h_j * g_{ip_i}, h_j * h_{iq_i})$. We denote these vectors as, $a_i = (h_{j*g_{ip_i}})$ and $b = (h_j * h_{iq_i})$. Then, we verify the condition $H(a-b)^T = 0$, with $det(H) = \pm 1$, such that the Hilbert basis is unimodular, if this condition is fulfil, we have finished the proof, if the ideal given by, $I_H =< (W_i^{h_j})^{g_{ip_i}} - (W_i^{h_j})^{h_{iq_i}} >=< W^a - W^b >$ not fulfill this condition, we apply newly the process of Hilbert basis, for the Hironaka's theorem , see [1] this guarantee, this process is finite. Thus always we can obtaint Hilbert basis to build a toric ideal $I_H$ to emmbed an S-system equation. **q.e.d.**
