## Supplementary material for "Topological Environment in Genetic and Metabolic Networks": Latex manuscript: interacttfpsample.bib

@MISC{ACSPD,
  title = {ACS Publications Division Home Page},
  url = {http://pubs.acs.org},
  lastchecked = {Nov~7, 2004}}

@MANUAL{USDA00,
  author = {{U.S. Department of Agriculture}},
  title = {Agriculture Fact Book 2000},
  organization = {U.S. Government Printing Office},
  address = {Washington, DC},
  year = {2000}}

@INCOLLECTION{Alm96,
  author = {Almlof, J. and Gropen, O.},
  title = {Relativistic Effects in Chemistry},
  booktitle = {Reviews in Computational Chemistry},
  editor = {Lipkowitz, K. B. and Boyd, D. B.},
  publisher = {VCH},
  address = {New York},
  year = {1996},
  volume = {8},
  pages = {206--210}}

@BOOK{ACS06,
  author = {{American Chemical Society, Committee on Analytical Reagents}},
  title = {Reagent Chemicals: Specifications and Procedures},
  edition = {10},
  address = {Washington, DC},
  year = {2006}}

@BOOK{Ban75,
  editor = {Bandy, A. R.},
  title = {The Chemistry of the Atmosphere: Oxidants and Oxidation in the Earth's Atmosphere},
  publisher = {Royal Society of Chemistry},
  address = {Cambridge, UK},
  year = {1995}}

@MASTERSTHESIS{Bla04,
  author = {Bl{\"a}ttler, T. M.},
  title = {Covalent Immobilization of Poly(\textsc{l}-lysine)-\textit{g}-poly(ethylene glycol) onto Aldehyde Plasma Polymer Coated Surfaces},
  type = {Diploma Thesis},
  school = {University of South Australia},
  year = {2004}}

@ARTICLE{Bro45,
  author = {Brosset, C.},
  title = {},
  journal = {Ark. Kemi, Mineral. Geol.},
  year = {1945},
  volume = {20A},
  pages = {No.~7}}

@ARTICLE{Car01,
  author =  {Caruso, R. A. and Susha, A. and Caruso, F.},
  title =   {Multilayered Titania, Silica and Laponite Nanoparticle Coatings on Polystyrene Colloidal Templates},
  journal = {Chem. Mater.},
  year =    {2001},
  volume =  {13},
  pages =   {400--409}}

@PHDTHESIS{Cha94,
  author = {J. S. Chandrakanth},
  title = {Effects of Ozone on the Colloidal Stability of Particles Coated with Natural Organic Matter},
  type = {Ph.D. Dissertation},
  school = {University of Colorado},
  address = {Boulder, CO},
  year = {1994}}

@UNPUBLISHED{Cha04,
  author = {K. Chatterjee and A. Visconti and C. J. Mirocha},
  title = {Deepoxy T-2 Tetraol: A Metabolite of T-2 Toxin Found in Cow Urine},
  note = {\emph{J. Agric. Food Chem.}, submitted for publication, 2004}}

@ARTICLE{Cha91,
  author = {Y. Chauvin and B. Gilbert and I. Guibard},
  journal = {Vib. Spectrosc.},
  year = {1991},
  volume = {1},
  pages = {299--304}}

@ARTICLE{Cha90,
  author = {Y. Chauvin and B. Gilbert and I. Guibard},
  journal = {J. Chem. Soc., Chem. Commun.},
  year = {1990},
  pages = {1715--1716}}

@PRESENTATION{Diz04,
  author = {B. Dizman and M. O. Elasri and L. J. Mathias},
  title = {Presented at the 227th National Meeting of the American Chemical Society},
  address = {Anaheim, CA},
  month = {March~28--April~1},
  year = {2004},
  note = {Paper POLY 229}}

@PHDTHESIS{Eri04,
  author = {T. A. Erickson},
  title = {Development and Application of Geostatistical Methods to Modeling Spatial Variation in Snowpack Properties, Front Range, Colorado},
  type = {Ph.D. Dissertation},
  school = {University of Colorado},
  address = {Boulder, CO},
  year = {2004}}

@INPROCEEDINGS{Gar88,
  author = {Garrone, E. and Ugliengo, P.},
  title = {},
  booktitle = {Structure and Reactivity of Surfaces, \emph{Proceedings of the European Conference}},
  address = {Trieste, Italy},
  month = sep # "~13--20, 1988",
  editor = {Zecchina, A. and Cost, G. and Morterra, C.},
  publisher = {Elsevier: Amsterdam},
  year = {1988}}

@BOOK{Gou02,
  author = {Gould, S. J.},
  title = {The Structure of Evolutionary Theory},
  publisher = {Belknap Press},
  address = {Cambridge, MA},
  year = {2002}}

@INCOLLECTION{Hil90,
  author = {Hillman, L. W.},
  title = {},
  booktitle = {Dye Laser Principles with Applications},
  editor = {Duarte, F. J. and Hillman, L. W.},
  publisher = {Academic},
  address = {New York},
  year = {1990},
  chapter = {2}}

@INCOLLECTION{Hol05,
  author = {Holbrey, J. D. and Chen, J. and Turner, M. B. and Swatloski, R. P. and Spear, S. K. and Rogers, R. D.},
  title = {Applying Ionic Liquids for Controlled Processing of Polymer Materials},
  booktitle = {Ionic Liquids in Polymer Systems: Solvents, Additives, and Novel Applications},
  editor = {Brazel, C. S. and Rogers, R. D.},
  series = {ACS Symposium Series},
  number = {913},
  publisher = {American Chemical Society},
  address = {Washington, DC},
  year = {2005},
  pages = {71--88}}

@BOOK{Kli04,
  author = {Kline, R. B.},
  title = {Principles and Practice of Structural Equation Modeling},
  edition = {Second},
  publisher = {Guilford Press},
  address = {New York},
  year = {2004}}

@ARTICLE{Kli05,
  author =  {Klingler, J.},
  title =   {Influence of Pretreatment on Sodium Powder},
  journal = {Chem. Mater.},
  year =    {2005},
  volume =  {17},
  pages =   {2755--2768}}

@INBOOK{LeC03,
  author = {Le Couteur, P. and Burreson, J.},
  year = {2003},
  title = {Napoleon's Buttons: How 17 Molecules Changed History},
  edition = {},
  address = {New York},
  publisher = {Jeremy P. Tarcher/Putnam},
  pages = {32--47}}

@MISC{Len04,
  author = {Lenssen, K. C. and Jantscheff, P. and Kiedrowski, G. and Massing, U.},
  title = {Cationic Lipids with Serine Backbone for Transfecting Biological Molecules},
  note = {Eur. Pat. Appl. 1457483, 2004}}

@BOOK{Leo05,
  editor = {S. R. Leone and A. E. McDermott and A. Paul},
  title = {Annual Review of Physical Chemistry},
  publisher = {Annual Reviews},
  address = {Palo Alto, CA},
  year = {2005},
  volume = {56}}

@ARTICLE{LP02,
  author = {Y. Lu and J. J. Pignatello},
  title = {},
  journal = {Environ. Sci. Technol.},
  year = {2002},
  volume = {36},
  pages = {4553--4561}}

@ARTICLE{LP04,
  author = {Y. Lu and J. J. Pignatello},
  title = {},
  journal = {Environ. Sci. Technol.},
  year = {2004},
  volume = {38},
  pages = {5853--5862}}

@PHDTHESIS{Mae04,
  author = {M{\"a}ckel, H.},
  title = {Capturing the Spectra of Silicon Solar Cells},
  school = {The Australian National University},
  year = {December 2004}}

@BOOK{McG02,
  title = {McGraw-Hill Encyclopedia of Science and Technology},
  edition = 9,
  publisher = {McGraw-Hill},
  address = {New York},
  year = {2002},
  note = {20 vols}}

@PHDTHESIS{Moh04,
  author = {M. Mohamed},
  title = {Waterjet Cutting Up to 900~MPa},
  school = {University of Hannover},
  address = {Germany},
  year = {2004}}

@INBOOK{Mor03,
  author = {Morris, R.},
  year = {2003},
  title = {The Last Sorcerers: The Path from Alchemy to the Periodic Table},
  edition = {},
  address = {Washington, DC},
  publisher = {Joseph Henry Press},
  pages = {145--158}}

@ARTICLE{Mul05,
  author = {Mullin, R.},
  title = {},
  journal = {Chem. Eng. News},
  year = {2005},
  volume = {83},
  number = {Oct~17},
  pages = {7}}

@MISC{Pet99,
  author = {Petrovick, P. R. and Carlini, E.},
  title = {Antiulcerogenic Preparation from \textit{Maytenus ilicifolia} and Obtaintion Process},
  note = {Br. Patent PI 994502, March~6, 1999}}

@INCOLLECTION{Pul04,
  author = {Puls, J. and Saake, B.},
  title = {Industrially Isolated Hemicelluloses},
  booktitle = {Hemicelluloses: Science and Technology},
  editor = {Gatenholm, P. and Tenkanen, M.},
  series = {ACS Symposium Series},
  number = {864},
  publisher = {American Chemical Society},
  address = {Washington, DC},
  year = {2004},
  pages = {24--37}}

@ARTICLE{Pus05,
  author = {Puskas, J. E. and Chan, S. W. P. and McAuley, K. B. and Shaikh, S. and Kaszas, G.},
  title = {},
  journal = {J. Polym. Sci., Part A: Polym. Chem.},
  year = {2005},
  volume = {43},
  pages = {5394--5413 and references therein}}

@BOOK{Sah04,
  editor = {Saha, B. C. and Hayashi, K.},
  title = {Lignocellulose Biodegradation},
  series = {ACS Symposium Series},
  number = {889},
  publisher = {American Chemical Society},
  address = {Washington, DC},
  year = {2004}}

@MISC{She04,
  author = {Sheem, S. K.},
  title = {Low-Cost Fiber Optic Pressure Sensor},
  note = {US Patent 6,738,537, May~18, 2004}}

@ARTICLE{Squ05,
  author = {Squires, S.},
  title = {Falling Short on Nutrients},
  journal = {The Washington Post},
  volume = {},
  month = oct # "~4",
  year = {2005},
  pages = {H1}}

@BOOK{Stoc01,
  editor = {J. H. Stocker},
  title = {Chemistry and Science Fiction},
  publisher = {American Chemical Society},
  year = {1998},
  address = {Washington, DC}}

@ARTICLE{Tay05,
  author = {Taylor, C. W. and Kumar, S.},
  title = {},
  journal = {Eur. J. Cancer},
  year = {2005},
  volume = {40},
  number = {Suppl.~1},
  pages = {781}}

@TECHREPORT{Tsc04,
  author = {Tschantz, B. A. and Moran, B. M.},
  title = {Modeling of the Hydrologic Transport of Mercury in the {UEFPC} Watershed},
  institution = {Lockheed Martin Energy Systems},
  address = {Bethesda, MD},
  month = {September},
  year = {2004}}

@INCOLLECTION{Wib74,
  author = {Wiberg, K.},
  title = {},
  booktitle = {Investigations of Rates and Mechanisms of Reactions},
  editor = {E. S. Lewis},
  series = {Techniques of Chemistry},
  volume = {VI, Part~I},
  publisher = {Wiley},
  address = {New York},
  year = {1974},
  pages = {764}}

@BOOK{WHO02,
  author = {{World Health Organization}},
  title = {Pathology and Genetics of Tumours of the Head and Neck},
  address = {Albany, NY},
  year = {2002},
  volume = {9}}

@PRESENTATION{Zie03,
  author = {K. D. Zientek and J. R. Eyler},
  title = {Presented at the 51st ASMS Conference on Mass Spectrometry and Allied Topics},
  address = {Montreal, Canada},
  month = jun # "~8--12, 2003"}
