## Supplementary material for "Topological Environment in Genetic and Metabolic Networks": Latex manuscript: interacttfpsample.tex

% interacttfpsample.tex
% v1.01 - June 2016

\documentclass[]{interact}
\usepackage{amsthm}
\usepackage{epstopdf}% To incorporate .eps illustrations using PDFLaTeX, etc.
\usepackage{subfigure}% Support for small, `sub' figures and tables

\usepackage[numbers,sort&compress,merge]{natbib}% Citation support using natbib.sty
\bibpunct[, ]{(}{)}{,}{n}{,}{,}% Citation support using natbib.sty
\renewcommand\bibfont{\fontsize{10}{12}\selectfont}% Bibliography support using natbib.sty
\renewcommand\citenumfont[1]{\textit{#1}}% Citation numbers in italic font using natbib.sty
\renewcommand\bibnumfmt[1]{(#1)}% Parentheses enclose ref. numbers in list using natbib.sty

\theoremstyle{plain}% Theorem-like structures
\newtheorem{theorem}{Theorem}[section]
\newtheorem{lemma}[theorem]{Lemma}
\newtheorem{corollary}[theorem]{Corollary}
\newtheorem{proposition}[theorem]{Proposition}

\theoremstyle{definition}
\newtheorem{definition}[theorem]{Definition}
\newtheorem{example}[theorem]{Example}

\theoremstyle{remark}
\newtheorem{remark}{Remark}
\newtheorem{notation}{Notation}

\begin{document}

%\articletype{Journal of Biological Dynamics}

\title{Topological Environment in Genetic and Metabolic Networks}

\author{
\name{Marco Polo Castillo-Villalba\textsuperscript{a}\thanks{CONTACT Marco Polo Castillo-Villalba.} }
\affil{\textsuperscript{a}Programa de Genómica Computacional, Centro de Ciencias Genómicas, Campus\\ Morelos, Universidad Nacional Autónoma de México, Av Universidad s/n, Col. \\Chamilpa, Cuernavaca, 62210, Morelos, México.}
}
\maketitle

\begin{abstract}

The analysis of large gene and metabolic networks is often hindered by unknown biochemical parameters and the nonlinear nature of classical S-system models. To address this, we introduce a framework based on combinatorial toric geometry computed with tools such as Normaliz, SageMath, it is worth mentioning this technique in not restrictive to integer vectors, there exists a natural extension to real geometries. Unlike traditional approaches, which rely on parameter dependent fixed points, our method constructs a Topological Environment derived from the dual space of kinetic orders, leading to what we call orthogonal enzyme kinetics. Within this topological setting, fixed points are computed on the algebraic torus, enabling the transformation of nonlinear dynamics into linear forms. Importantly, these fixed points are independent of kinetic parameters and depend only on network topology and interaction signs. Applying this methodology to gene circuits involved in circadian rhythms, we reproduce previously reported oscillatory physiologies.

%Analysis of large molecular and gene networks, where many biochemical parameters are unknown, can be addressed within the framework of \textbf{combinatorial toric geometry}.
%Our approach involves analyzing gene networks using \textbf{S-systems} (a power-law formalism of enzyme kinetics) developed by M.A. Savageau.
%One of the primary challenges in solving these models is their highly non-linear nature, compounded by the frequent lack of known kinetic parameters. A well established technique to address this is to transform the system into a \textbf{linear dynamical system} by analyzing its fixed points, which are computed in steady state. However, these fixed points themselves typically depend on unknown parameters. In this work, we propose a novel approach in which we redefine the model in a new coordinate system using a linear transformation calculated from specialized tool such as \texttt{Normaliz, SageMath and Singular}, it is worth mentioning this technique in not restrictive to integer vectors, there exists a natural extension to rational or real geometries cones. As a natural consequence of this geometric framework, we define the concept of a \textbf{Topological Environment} for gene networks. This environment is constructed from the dual space of kinetic orders, yielding what we refer to as \textbf{orthogonal enzyme kinetics}. Within this topological space, we compute \textbf{fixed points on the torus}, enabling us to transform the original nonlinear model into a linear form within this ambient space. The key result of our work is that these fixed points are \textbf{independent of kinetic parameters} and instead depend solely on the \textbf{topology of the network} and the \textbf{signs of gene interactions}. Based on this methodological approach, was applied in a gene and metabolic circuit, involved in natural circadian rythms reproducing different oscillatory phisiologies, previously reported in the literature.

\end{abstract}

\begin{keywords}
Power-Law Formalism, S-Systems, Fix points on torus, Quantitative phenotype; Oscillatory phenotypes; Dominant fluxes
\end{keywords}


\section{Introduction}

Mathematical models are essential tools for analyzing and understanding the behavior of genetic and metabolic networks. In such models, the \textbf{vertices} typically represent molecular species such as genes, mRNAs, proteins, or metabolites while the \textbf{edges} correspond to the regulatory interactions among these components.

Although a substantial amount of \textbf{qualitative data} is available, enabling the inference of biological network topology, information about the \textbf{kinetic parameters} governing the underlying biochemical reactions remains relatively scarce. This lack of quantitative data presents a significant challenge in constructing and analyzing large-scale biological networks.

%Nonetheless, several mathematical techniques exist to approximate the parameter space, thereby facilitating the study of the \textbf{dynamical properties} of complex biological systems.

Nonetheless, several mathematical techniques exist to approximate the parameter space, thereby facilitating the study of the \textbf{dynamical properties} of complex biological systems. In the context of statistical learning machines and learning manifolds, a considerable number of novel works have been developed in recent years for the analysis of large-scale networks 
(see \cite{rev21}, \cite{rev22}, \cite{rev23}, \cite{rev24}, \cite{rev25}). These computational methodologies have a strong foundation in Information Geometry, which is based on Differential Geometry and Parametric Multivariate Statistics. The main challenge observed in these methodologies concerns the identifiability of the statistical machines in their parametric space. Most of these techniques correspond to non-identifiable models, which have been extensively studied (see \cite{Watanabe1}). The problem of non-identifiable models is related to the existence of singularities in high-dimensional parameter spaces, and the performance of learning from large data sets strongly depends on these singularities. Such models are often referred to as \emph{Singular Learning Machines}. The techniques of toric geometry and the topological environment may serve as complementary approaches to these computational models, as discussed in the conclusion of this work.\\

There are numerous formalisms available for modeling gene networks. Among them, the \textbf{Boolean formalism} developed by R. Thomas \cite{Thomas} focuses primarily on gene regulation and typically does not include metabolic reactions. Another widely used approach is \textbf{Flux Balance Analysis (FBA)}, developed by B.O. Palsson \cite{BOPalsson1,BOPalsson2,BOPalsson3}, which models metabolism based on the stoichiometric matrix but has limited capacity to incorporate genetic regulatory mechanisms. In addition, stochastic methods for the simulation of the entire cell in E. coli, \cite{Multiomics}, are of consideration, with the finality of predicting genome-wide biochemical concentrations and growth dynamics for E. coli, using a large number of sophisticated computational techniques (neural networks, optimization, and NLP).\\

In contrast, modeling the parameter space using \textbf{S-systems}, as recently in \cite{bib54} and previous works \cite{MASavageau1976}, \cite{SavageauCoelhoFasaniSalvador2009}, \cite{MASavageauVoit1987}, \cite{MASavageau2001}, \cite{SavageauOptimal}, \cite{rev13}, \cite{LomnitzSavageau2013}, \cite{LomnitzSavageau2014}, \cite{LomnitzSavageau2015}, \cite{LomnitzSavageau2016} allows for the integration of both gene regulation and biochemical reactions within a unified mathematical framework. Also there are many works developed about this formalism, with interesting implications in biology (see \cite{rev11}, \cite{rev12}, \cite{rev13}, \cite{rev14}, \cite{rev15}), in some cases these methodologies open deep questions related with the dynamical analysis in metabolic network, mainly associated to stable quantitative phenotypes and the existence of hidden oscillatory behaviours, such as is interest for this work, the conjeture in Voit. E. O. \cite{rev15}, about an S-system in its non linear canonical representation with the monomial matrix transformation for matrices $G$ $H$, where $\dot{X} = X^G - X^{H}$,   when the $det(G-H)=0$ the conditions of stability are not clear by the central manifold theorem, and less the geometries or manifolds corresponding to this condition, for the cases with $det(G-H) \neq 0$, are well known the associated dynamics. In the \textbf{Theorem 3.3}, the author elucidates a perspective to face this challenge, also the commutativity of a diagram of maps (lattice cones, formal polynomials and algebriac varieties) is given in support information that shows that classical fix points in dynamical systems are contained in the fix points on torus, such that, the fix points are goemetries ruled by objects more general such as, binomial ideals or generator of binomials. Also is of interest for the author, the interesting conjetures and results addressed in the analysis provided, \cite{rev12}. Opening possiblities to formalize the relationship between genotype-phenotype. Finding an one equilibrium point between the space of stoichimoetric parameters from balance of general action mass into the kinetic order space defined by signed lattice cones, this fact is deeply conected through the monomial map defined in toric geometry, also is addressed in the disccusion of this work. Finally, the toric characterization in the work \cite{rev14} has a interesting treatment from point of view of steady-state for algebraic biochemical reactions, its methodology based in Grobner bases for the coeficient spaca is valuable to gain knowledge about the topological hidden properties in molecular networks. In contrast with this work, through of an analysis with Hilbert basis into the space of kinetic orders, a local dynamical analysis is possible in gene regualtion and metabolism has been gained in this methodology.\\   

Although these approaches are highly effective, our intention is not to compete with them. In contrast, our goal is to contribute to this growing set of modern modeling techniques by incorporating \textbf{computational algebraic geometry}, \cite{DCoxLittleOshea2004}, \cite{DCoxLittleSchenck2010},\cite{BSturmfels1996}, \cite{Normaliz}. In this context, the proposed concept of the \textbf{ topological environment} should be viewed as a tool to extract hidden topological features in large-scale networks, or as a complementary method in scenarios where detailed parameter information is not available.\\


In the present work, we model genetic and metabolic networks using the \textbf{Power-Law Formalism} and \textbf{S-systems}. As mentioned above, these models were originally proposed and extensively studied by M.A. Savageau \cite{MASavageau1976, MASavageauVoit1987, MASavageau2001}. Our goal is to contribute to the extension of these models from small scale systems to large-scale genetic and metabolic networks.\\

The primary technique used thus far in the literature is \textbf{ non-linear dynamical analysis}, which involves linearizing the system around its \textbf{fixed points}, calculated in steady state. Once linearized, the system can be solved analytically through the computation of eigenvalues of the associated state matrix. However, in the context of gene networks, this approach is more complex, as the fixed points required for linearization also depend on \textbf{unknown kinetic orders and parameters}.\\

The central aim of this work is to provide a \textbf{geometric approach} for describing fixed points in a dual space, referred to as the \textbf{dual cone}, where these points are independent of kinetic orders. In this work, the S-systems can be visualized as \textbf{toric varieties} \cite{MPCastilloLauraJulio2017}, a structural property that has also been observed by other authors in the modeling of biochemical reaction networks \cite{MASavageau1976, MASavageauVoit1987, MASavageau2001}.

Building upon these results, we introduce the concept of the \textbf{Topological Environment}, where analysis is localized to a single element in the network, keeping all other elements fixed i.e., focusing on the \textit{topological neighborhood} of the chosen species. To construct this framework, we define a dual space called \textbf{orthogonal enzyme kinetics}, which is obtained by transforming the S-system equations into \textbf{toric coordinates} through a \textbf{monomial transformation}. This process uses tools from computational algebraic geometry, such as the computation of the \textbf{Hilbert basis for monoids}, and is implemented using software packages like \texttt{Normaliz or Singular} \cite{Normaliz, DGPS}.\\

In this transformed ambient space, we compute a special class of fixed points known as \textbf{fixed points on the algebraic torus}, \cite{GEwald1993}, \cite{Brasselet}. Crucially, we show that these fixed points \textbf{do not depend on kinetic parameters}. Furthermore, we can linearize the S-system equations using the \textbf{Jacobian matrix} technique, thereby obtaining a linear system of ordinary differential equations that governs the dynamics of the metabolite or species of interest. The solutions of these linearized equations allow us to describe the system’s behavior in the original variable space.\\

This work is organized into the following sections:

In \textbf{Section 2}, we review the theoretical concepts related to the modeling of biochemical networks using \textbf{S-systems}, and introduce key ideas from \textbf{toric algebraic geometry} and their application to S-system equations. The section concludes with a discussion on the importance of \textbf{fixed points on the torus}, which are used to derive constraints on kinetic orders and to demonstrate the linearization of S-systems through a worked example.

In \textbf{Section 3}, and specifically in Subsection 3.1, we formally define the concept of the \textbf{Topological Environment} in genetic and metabolic networks. We provide a theoretical example to aid in the understanding of this concept, which is closely related to the idea of \textbf{orthogonal enzyme kinetics}. Additionally, we introduce a related concept that we refer to as \textbf{environment circuits}.

In \textbf{Section 4}, we develop an example and present results based on a widely studied system by M.A. Savageau \cite{LomnitzSavageau2013}: an \textbf{oscillatory synthetic gene circuit} regulated under different control modes in accordance with \textbf{Demand Theory} \cite{MASavageauVoitIrvine987}. This example gives rise to distinct regions in the kinetic parameter space, which are associated with \textbf{oscillatory quantitative phenotypes} sustained oscillatory behaviors characterized in logarithmic space according to metabolite concentration. We reproduce the dynamics of these phenotypes involved in natural circadian rythms within the Topological Environment. It also has been included a toy example applying the methodology, in a final subsection with the aim to clear concepts. The complete analysis was included in the support information.

Finally, in the \textbf{Discussion}, we outline potential future applications of the Topological Environment framework to the study of large-scale genetic and metabolic networks.


%%%%%%%%%%%%%%%%%%%%%%%%%%%%%%%%%%%%%%%%%%%%%%%%%%%%%%%%%%%%%%%%%%%%%%%%%%%%%
%%%%%%%%%%%%%%%%%%%%%%%%nueva secccion%%%%%%%%%%%%%%%%%%%%%%%%%%%%%%%%%%%%%%%%%%%

\section{Methods and Definitions}

\subsection{Biochemical networks and applications of Toric Algebraic Geometry.}

In this section, we introduce biochemical systems specifically genetic and metabolic networks modeled using the power-law formalism of enzyme kinetics, as studied and developed by M. Savageau \cite{MASavageau1976}, \cite{MASavageauVoit1987}, \cite{MASavageau2001}. In previous work, we demonstrated that these systems can be represented as toric varieties.\\


\textbf{Michaelis-Menten Kinetics}.\\

We present the classical form of the Michaelis–Menten rate law as a representative example, without loss of generality. It is well known that biomass balance in biochemical reactions can be described by more complex expressions; see Alberty \cite{Alberty}, for further discussion.\\

\begin{equation}
\dot{X}_{i}= \dfrac{X_{max}[S]}{K_{m}+ [S]}.
\end{equation}

Here, $\dot{X}_{i} $, represents the biochemical change in concentration of the $ i- $esim species of the cascade of reaction networks. $ X_{max} $ represents the maximum ratio achieved for the system, and $K_{m}  $ is the Michelis constant that depends on kinetic parameters, and $ [S] $ the substrate concentration. In complex reactions, in general, the expressions for $ X_{max} $ and $K_{m}  $ can be more complicated.\\

In the theory of S-systems, the rate law for the expression of $\dot{X}_{i}  $, is analyzed in a representation of Taylor's expansion, i.e., in a formal analytical treatment, as follows:\\

\begin{equation}
\dot{X}_{i}=\sum^{k}_{j=1} a_{ij}  X^{n}_{ij}.
\end{equation}

\textbf{Formalism of power law for biochemical systems.}\\\\

The power law formalism is a particular representation of enzyme kinetics, formally Taylor's expansion in several variables, which is capable of describing biochemical reactions of metabolic and gene regulatory mechanisms, formulated as rational functions, produced by the balance of mass equations (GMA).

\begin{equation}
 \dot{X}_{i}= \sum_{k=1}^{p_{i}} \alpha_{ik} \prod_{j=1}^{m}X_{j}^{g_{ijk}} - \sum_{k=1}^{q_{i}} \beta_{ik}\prod_{j=1}^{m} X_{j}^{h_{ijk}},\: i=1,...,n.
\end{equation}

This expression represents the basic principle of mass conservation, called \textbf{general mass action (GMA)}, in the form of the power law formalism. Each term is associated with elements of a network of reactions, where $\dot{X}_{i} $ represents a derivative of $ X_{i} $ over time t. This expression represents the rate law of degradation or production of any $ i $-esim specie $ X_{i} $ (gene, mRNA, proteins or metabolites) in the network. This variation in time describes the flux of biomass through all elements in any biochemical reaction and genetic network. The first sum is that of the products that contribute to the production of the species $ X_{i} $ and the terms of the second sum are elements of consumption including degradation or dilution of $ X_{i} $, \cite{MASavageau1976}, \cite{MASavageau2001}.\\

\begin{definition}\label{Definition 3.1} (\textbf{Dominant terms}). A dominant term is defined as the largest term of a given sign for an equation of the GMA-system (equation defined above, General Mass Action); the dominant terms with positive and negative signs are the \textbf{dominant positive term} and the \textbf{dominant negative term}, respectively, see ref. \cite{MASavageau1976}.\end{definition}\

It is possible to experimentally detect dominant fluxes in a metabolic reaction network, as the metabolic pathway with the highest amount of expression of the enzymes involved, measured by GFP.\\

We will see later, in section 4., that one way to analyze dominant fluxes is equivalent to study main branches of the S-system equations with toric algebraic geometry, and we give a geometrical technique to compute them.\\

%\textbf{Definition 6.0. (Dominant terms).} For the expression defined above, for all $ i=1,...,n $, the largest term in the sum is defined as the dominant term; if this term corresponds to a term in the positive sign sum, it is defined as the dominant positive term, and similarly for the negative sum, it is defined as the dominant negative term.\\

\begin{definition}\label{Definition 3.2} (\textbf{S-systems}). Using the concept defined above, we can have a finite number of combinations of dominant terms, exactly  \[ \prod_{j=1}^{m} p_{j}q_{j} ; \]  partitions of the space of concentrations $(X_{1},...,X_{n}) $, where each partition is a dominant sub-system and it is associated with a particular system of equations, called \textbf{S-systems}, \cite{MASavageauVoit1987}, \cite{MASavageau1976}, \cite{LomnitzSavageau2013}, \cite{LomnitzSavageau2014}, \cite{LomnitzSavageau2015}; as shown below
\begin{equation}
\dot{X}_{i}=  \alpha_{ip_{i}} \prod_{j=1}^{m} X_{j}^{g_{ip_{i}}} -  \beta_{iq_{i}} \prod_{j=1}^{m}X_{j}^{h_{iq_{i}}}, \, i=1,...,n.\\
%\[ 0=  \alpha_{ip_{t+1}} \prod_{j=1}^{m}X_{j}^{g_{ip_{t+1}}} -  \beta_{iq_{t+1}} \prod_{j=1}^{m}X_{j}^{h_{iq_{t+1}}}, \, t=0,...,m,
\end{equation}

for details, see M. Savageau \cite{MASavageauVoit1987}.\end{definition}\

Note in this last definition that the sums were neglected, in contrast to the power-law complete formalism. An S-System only consists of dominant terms, i.e., mathematical expressions of differential equations with only two monomial terms.\\
%%%%%%%%%%%%red bioquimica
\subsection{Toric Algebraic Geometry of Genetic and Metabolic Networks}

In this part of our work, we apply key concepts from algebraic geometry to analyze an auto-repressed gene circuit studied by Savageau \cite{MASavageau2001}. The analysis focuses on the effect of allolactose, the inducer of the Lac transcription factor, and its role in the allosteric transition of the Lac repressor bound to DNA into the Lac allolactose complex, which subsequently unbinds from the DNA. These types of regulatory reactions are common in microbial systems and are therefore considered fundamental design principles in genetic circuits.

\textbf{Objectives of Applying Computational Toric Geometry}

Our primary aim in this example is to demonstrate how a geometric approach can be effectively used to study concentration changes of any species in a gene network.

\begin{itemize}
  \item \textbf{First.} We construct new topological structures associated with gene and metabolic networks that do not depend on the order of the kinetic parameters. These structures include: \textbf{fixed points on the algebraic torus}.
  
  \item \textbf{Second.} These fixed points depend only on the signs of species interactions and the topology of the network.
  
  \item \textbf{Third.} It is sufficient to analyze a subset of the S-system equations to study a specific element or metabolite of interest. There is no need to solve the full set of S-systems to determine the quantitative phenotypes associated with that metabolite.
  
  \item \textbf{Fourth.} This method is applied specifically to the S-system equation corresponding to the molecule or metabolite under investigation.
  
  \item \textbf{Fifth.} We transform non linear algebraic expressions into linear forms within an appropriate geometric framework namely, a toric variety.
  
  \item \textbf{Sixth.} These new linear algebraic expressions are derived from a set of ordinary differential equations. Their solutions describe the local dynamics of the metabolite or molecule of interest without requiring the solution of the entire system of S-equations.\\
\end{itemize}
\\

\textbf{Gene Network of a Regulator and Inducer as a Basic Design Principle in Gene Circuits.}\\
\\
%We apply this methodology to the case of an allosteric system by end-product inhibition for a pathway of length $n=3$, which exhibits oscillatory behaviour. This system models a biosynthetic pathway, such as the biosynthesis of homoserine regulated by methionine (see EcoCyc pathways). In the following, we provide conditions in the topological environment for the nonlinear expression of $X_{1}$. Its local dynamics confirm this quantitative phenotype, (see Fig.~1). Here, the allosteric protein $X_{1}$ catalyzes the enzyme $X_{2}$, after which the enzyme $X_{3}$ antagonizes the activity of the allosteric protein through eight regulatory subunits ($g=8$) in its quaternary structure (orange points), as shown in \cite{SavageauOptimal}. A detailed version of this example is provided in the Supporting Information.\\


We now consider the set of equations of the S-system associated with the genetic circuit shown in Figure~1, and analyze the concentration changes that govern the production of the allolactose inducer, denoted by the variable \( X_3 \).

In simple terms, the procedure for writing S-system equations involves representing the network as a cascade of information. Alternatively, when focusing on a specific node in the network, the incoming edges correspond to positive terms (i.e., production), while the outgoing edges represent negative terms (i.e., degradation) in the S-system representation.


%%%%%%%%%%%%%%%%%%%%%%%%%%%%%%%%%%%%%%%%%%%%%%%%%%%%%%%%%%%%%%%

%%%%%%%%%%%%%%%%%%%%%%%%%%%%%%%%%%%%%%%%%%%%%%%%%%%%%%%%%%%%%%
\begin{figure}[H]
\centering
\makebox{\includegraphics[height=1.35in]{Circuit1dpi.png}}
\caption{\label{fig_simvarex} Schematic representation of an oscillatory gene circuit with six elements, (M.A. Savageau) \cite{MASavageau2001}.}
\end{figure}

\begin{equation}
 \dot{X}_{1}= \alpha_{3,5,6} X^{g1,3}_{3} X^{g1,5}_{5} X^{g1,6}_{6} - \beta_{1} X^{h1,1}_{1},\\
\end{equation}
\begin{equation}
 \dot{X}_{2}= \alpha_{1,7} X^{g2,1}_{1} X^{g2,7}_{7} - \beta_{2} X^{h2,2}_{2},\\
\end{equation}
\begin{equation} 
 \dot{X}_{3}= \alpha_{2,8} X^{g3,2}_{2} X^{g3,8}_{8} - \beta_{3,2} X^{h3,2}_{2} X^{h3,3}_{3},\\
\end{equation}
\begin{equation} 
 \dot{X}_{4}= \alpha_{3,5,6} X^{g4,3}_{3} X^{g4,8}_{5} X^{g4,6}_{6} - \beta_{4} X^{h4,4}_{4},\\
\end{equation}
\begin{equation} 
 \dot{X}_{5}= \alpha_{4,7} X^{g5,4}_{4} X^{g5,7}_{7} - \beta_{5} X^{h5,5}_{5}.\\
\end{equation}

Let us now proceed to select the S-system equation corresponding to the inducer, \( \dot{X}_{3} \). To construct the set of terms associated with the polynomial \( \dot{X}_{3} \), we extract the exponents of each monomial term in this case, there are only two terms. These exponents are then expressed in vector notation as follows. This collection of exponent vectors is referred to as the \textbf{support of the polynomial}; for further details, see \cite{DCoxLittleOshea2004}, \cite{DCoxLittleSchenck2010}, \cite{GEwald1993}, \cite{Brasselet}.


\begin{center}
$  Supp( \dot{X}_{3})= \left\lbrace (0,g_{32},0,0,0,0,0,g_{38}),\, (0,h_{32},h_{33},0,0,0,0,0) \right\rbrace $.
\end{center}

In this work, we exploit all the major topological information, with the purpose of identifying new topological invariant in gene networks, as we can see later.\\

Next, we express the support associated with the polynomial in matrix notation by introducing a new vector obtained as the difference between the two original exponent vectors, namely:
\[
(0, g_{32}, 0, 0, 0, 0, 0, g_{38}) - (0, h_{32}, h_{33}, 0, 0, 0, 0, 0).
\]
The purpose of this difference vector is to construct a closed geometric object known as a \textbf{cone}, or more specifically, the \textbf{convex hull} of the support set. For formal definitions and geometric background, see \cite{GEwald1993}.\\

If the support contains more than two vectors, we compute pairwise differences and retain only the two extreme vectors, selected according to the lexicographic order technique; see \cite{DCoxLittleOshea2004} for details.\\

The positions in the resulting row matrix correspond to the number of variables (or components) associated with the molecular species in the network. The resulting matrix is as follows:\\\\

\textbf{Cone associated to $ \dot{X}_{3} $}.\\\\


$\sigma_{\dot{X}_{3}}=      
\begin{pmatrix}
0&g_{32}&0&0&0&0&0&g_{38}\\
0&(g_{32}-h_{32})&-h_{33}&0&0&0&0&g_{38}\\
0&h_{32}&h_{33}&0&0&0&0&0\end{pmatrix}  
$, \\\\

Afterwards, we construct a new matrix from $\sigma_{\dot{X}_{3}}$, filling the rows of this matrix with orthogonal vectors to the vectors to $\sigma_{\dot{X}_{3}}  $, i.e., 

\begin{center}
$ <(0,g_{32},0,0,0,0,0,g_{38}),(1,0,1,1,1,1,1,0)> =0$, 
\end{center}

where $ < , > $, represents of usual scalar inner product. We fill for computational simplicity with $ 1 $ values this matrix, including any vector $ b $, keeping the constraint $ <(0,g_{32},g_{38},0,0,g_{16}, b > \geq 0 $ as valid. This matrix is called the \textbf{Dual-cone} $\sigma^{\vee}_{\dot{X}_{3}}  $ to the cone $ \sigma_{\dot{X}_{3}}  $, see formal definition in, \cite{GEwald1993}.\\

\textbf{Dual-cone associated to $ \dot{X}_{3} $}.\\\\

$\sigma^{\vee}_{\dot{X}_{3}}=      
\begin{pmatrix}
1&0&1&1&1&1&1&0\\
1&0&0&1&1&1&1&0\\
1&0&0&1&1&1&1&1\end{pmatrix}  
$.\\ \\

We observe that the vectors in both the support and the cones have integer entries $g_{ij}
$; these are known as \textbf{lattice vectors}, since they lie in the integer lattice \( \mathbb{Z}^{n} \), which possesses the structure of a lattice (see \cite{GEwald1993} for details).\\ 
It is important to remark at this point that this methodology is not restricted to the use of integer coordinates. For example, it is always possible to define a map from rational cones to lattice cones, and vice versa. That is, one can define \[\rho : \sigma \cap \mathbb{Q}^{n} \longmapsto \sigma \cap \mathbb{Z}^{n}\] by multiplying by the least common denominator. 
Given a rational vector \[v_{\mathbb{Q}} = \left(\tfrac{1}{4}, \tfrac{2}{5}, \tfrac{7}{13}, \tfrac{1}{7}\right) \in \sigma \cap \mathbb{Q}^{4},\] we obtain \[\rho(v_{\mathbb{Q}}) = 910 \cdot \left(\tfrac{1}{4}, \tfrac{2}{5}, \tfrac{7}{13}, \tfrac{1}{7}\right) = (455, 364, 980, 130) \in \sigma \cap \mathbb{Z}^{n}.\] Conversely, by dividing by the least common denominator, one can define 
\[\rho^{-1}: \sigma \cap \mathbb{Z}^{n} \longmapsto \sigma \cap \mathbb{Q}^{n} \subset \sigma \cap \mathbb{R}^{n}.\] In summary, given any rational cone \[C = \text{Cone}(v_{1}, \ldots, v_{n}) \subseteq \sigma \cap \mathbb{Q}^{n}, \quad v_{i} = \frac{u_{i}}{d_{i}},\] we take \[
D = \mathrm{lcm}(d_{1}, \ldots, d_{n}),
\] and multiply the whole cone to embed it into a lattice cone, \[DC = \sigma \cap \mathbb{Z}^{n},
\] see \cite[Chap.~V, p.~144]{GEwald1993}.\\


Next, we compute the intersection of the dual cone with the lattice \( \mathbb{Z}^{8} \) in order to endow the cone with additional algebraic properties. This intersection is denoted as: $ S_{\sigma} = \sigma^{\vee}_{\dot{X}_{3}} \cap \mathbb{Z}^{8}, $
where \( S_{\sigma} \) is referred to as a \textbf{monoid} or \textbf{semigroup}. According to \textbf{Gordan's Lemma} (see \cite{GEwald1993}), every monoid is finitely generated; that is, it admits a finite set of generators.\\

We compute these generators using the software \texttt{Singular} (available at: \url{https://www.singular.uni-kl.de:8003}) \cite{DGPS}. The resulting set of generators is known as the \textbf{Hilbert basis} \cite{BSturmfels1996}. In matrix notation, the Hilbert basis is given by:\\\\


%\begin{center}
$ H_{\dot{X}_{3}}=      
\begin{pmatrix}

1&0&-1&0&0&0&0&-1\\
0&0&1&0&0&0&0&0\\
0&0&0&0&0&0&0&1\\
0&1&0&0&0&0&0&0\\
-1&0&0&1&0&0&0&0\\
-1&0&0&0&1&0&0&0\\
-1&0&0&0&0&1&0&0\\
-1&0&0&0&0&0&1&0\\
0&-1&0&0&0&0&0&0\\
1&0&0&-1&0&0&0&0\\
1&0&0&0&-1&0&0&0\\
1&0&0&0&0&-1&0&0\\
1&0&0&0&0&0&-1&0\end{pmatrix}  $\\ \\
%\end{center}

We select a subset of rows from the Hilbert basis matrix in order to construct a new matrix with the same number of variables as the original polynomial. The only constraint for choosing these row vectors is that the resulting matrix must have a determinant equal to \( \pm 1 \).\\\\


%\begin{center}
$ [H_{\dot{X}_{3}}]_{8\times 8}=      
\begin{pmatrix}

1&0&-1&0&0&0&0&-1\\
0&-1&0&0&0&0&0&0\\
0&0&1&0&0&0&0&0\\
1&0&0&-1&0&0&0&0\\
1&0&0&0&-1&0&0&0\\
1&0&0&0&0&-1&0&0\\
1&0&0&0&0&0&-1&0\\
0&0&0&0&0&0&0&1\\
\end{pmatrix}  $\\ \\
%\end{center} 

We obtain the following matrix, which has determinant \( \pm 1 \); such a matrix is referred to as a \textbf{unimodular matrix}. The cone generated by these new vectors is known as a \textbf{regular cone}, a topological condition associated with the concept of completeness in algebraic geometry. For technical details, see \cite{GEwald1993}, \cite{DCoxLittleOshea2004}.\\

Next, we take the transpose of the matrix \( [H_{\dot{X}_{3}}]_{8\times 8} \) to construct a \textbf{monomial transformation} (or change of coordinates) for the equation \( \dot{X}_{3} \), as follows:\\\\


%\begin{center}
$ [H_{\dot{X}_{3}}]^{T}_{8\times 8}=      
\begin{pmatrix}

1&0&0&1&1&1&1&0\\
0&-1&0&0&0&0&0&0\\
-1&0&1&0&0&0&0&0\\
0&0&0&-1&0&0&0&0\\
0&0&0&0&-1&0&0&0\\
0&0&0&0&0&-1&0&0\\
0&0&0&0&0&0&-1&0\\
-1&0&0&0&0&0&0&1\\
\end{pmatrix}  $\\\\

%\end{center} 


We associate the rows of the Hilbert matrix with the exponents of monomials, thereby defining a change of coordinates known as a \textbf{monomial transformation}, or more formally, a homomorphism of semigroups into the algebraic torus
\[
u = (u_{1}, u_{2}, u_{3}, u_{4}, u_{5}, u_{6}, u_{7}, u_{8}) \in (\mathbb{C}^{*})^{8},
\]
as described in \cite{GEwald1993}. This transformation maps each integer vector \( a_i \) to a monomial \( X^{a_i} \), that is, \( a_i \longrightarrow X^{a_i} \).

The specific monomial coordinate transformations are given by:
\begin{align*}
X_{1}(u) &= u_{1} u_{4} u_{5} u_{6} u_{7}, \\
X_{2}(u) &= u_{2}^{-1}, \\
X_{3}(u) &= u_{1}^{-1} u_{3}, \\
X_{4}(u) &= u_{4}^{-1}, \\
X_{5}(u) &= u_{5}^{-1}, \\
X_{6}(u) &= u_{6}^{-1}, \\
X_{7}(u) &= u_{7}^{-1}, \\
X_{8}(u) &= u_{1}^{-1} u_{8}.
\end{align*}

We observe that the exponents of the new variables are determined by the rows of the Hilbert matrix. This transformation will be substituted into the original equation \( \dot{X}_{3} \) in the subsequent analysis.\\\\


\textbf{Toric variety}.\\

We define a \textbf{toric variety} as the image of a set of algebra homomorphisms, where each map is defined by the monomials computed above. We denote the variety by \( X_{\sigma} \), and the set of maps \( \varphi_i \) is given by:


%\begin{flushleft}
\begin{equation}
X_{\sigma}=\left\lbrace X_{1}(u), X_{2}(u), X_{3}(u), X_{4}(u), X_{5}(u), X_{6}(u) , X_{7}(u), X_{8}(u) \right\rbrace \,\subseteq\, \mathbb{C}^{*} .
\end{equation}

%Each map is defined as $h_{i}  $ take elements in $(\mathbb{C}^{*})^{8}$ and are mapped to algebraic torus $T= \mathbb{C}^{*} $, i.e., the multiplicative group of complex numbers. The torus action $T $ on a toric variety $  X_{\sigma} $, we write 
%$ T \times X_{\sigma} \,\longmapsto \,X_{\sigma} $, in our example, is given by the following expressions.\\
\textbf{Action on Torus.}\\

We define the \textbf{torus action} in our example, denoted by \( t \), using the same monomial maps as those defining the toric variety \( X_{\sigma} \subseteq (\mathbb{C}^{*})^{n} \). However, in this context, the maps are not considered as a set, but rather as \textbf{polynomial coordinate functions}. For technical details, see \cite{GEwald1993}.\\


%\begin{flushleft}
\begin{equation}
t=(X_{1}(u), X_{2}(u), X_{3}(u), X_{4}(u), X_{5}(u), X_{6}(u) , X_{7}(u), X_{8}(u))\,\in\,(\mathbb{C}^{*})^{8} .
\end{equation}
%\end{flushleft}

The action of \( t \) on the variety \( X_{\sigma} \) is defined by component-wise complex multiplication, that is, $ t*X_{\sigma}=(X_{1}(u)*x_{1}, X_{2}(u)*x_{2}, X_{3}(u)*x_{3}, X_{4}(u)*x_{4}, X_{5}(u)*x_{5}, X_{6}(u) *x_{6}, X_{7}(u)*x_{7}, X_{8}(u)*x_{8}) $, where $ (x_{1},x_{2},x_{3},x_{4},x_{5}, x_{6},x_{7},x_{8})\,\in\,  X_{\sigma}$.
This type of map generates new topological invariants in the network. Two important examples are the \textbf{orbits} of the torus action and the corresponding \textbf{fixed points}. In what follows, we focus specifically on the fixed points, as they are essential for analyzing the dynamics of gene and metabolic networks.\\

\textbf{Fix points}.\\

In this work, the \textbf{fix points} on torus \cite{Brasselet}, are an important topological property associated to genetic-metabolic networks. We offer a practical formula to compute them.\\

Given the lattice vector $ \lbrace b_{1},...,b_{n} \rbrace$ generate the cone $ \sigma $, now let us consider the elements of Hilbert Basis $\lbrace h_{1},...,h_{r} \rbrace$ which generate the monoid $ S_{\sigma^{\vee}} $, associated to the dual-cone $\sigma^{\vee}  $. The fix points on torus $ X_{\tau} $ are computed by the following limit of maps,\\\\


\begin{equation}
 X_{b_{i}} = \lim_{Z \longrightarrow 0} Z^{<b_{i},h_{j}>} =(\lim_{Z \longrightarrow 0}Z^{<b_{i},h_{1}>},...,\lim_{Z \longrightarrow 0}Z^{<b_{i},h_{r}>} ).
\end{equation}

Running $ j =1,..,r $, and $ i$ fixed index, with values in, $i=1,..,n $, and $ <, > $, is the inner product of vectors. We see, how we can calculate the fix points in our example, formula, see in \cite{Brasselet}.\\

For example we calculate the fix point using the formula above, we take the cone $ \sigma $ and the face generated by the vector ray,\\


\begin{center}
$\tau= (0,(g_{32}-h_{32}),h_{33},0,0,0,0,g_{38}) $. 
\end{center}

We first calculate the scalar product, $ <,> $, to this face, projecting over each generator from the Hilbert basis, as we can see, in the following,\\

\begin{flushleft}
$ \left\langle (0,(g_{32}-h_{32}),h_{33},0,0,0,0,g_{38}),\, (1,0,-1,0,0,0,0,-1)  \right\rangle = -(h_{33}+g_{38})$,
$ \left\langle (0,(g_{32}-h_{32}),h_{33},0,0,0,0,g_{38}),\, (0,-1,0,0,0,0,0,0)  \right\rangle = -(g_{32}-h_{32}) $,
$ \left\langle (0,(g_{32}-h_{32}),h_{33},0,0,0,0,g_{38}),\, (0,0,1,0,0,0,0,0) \right\rangle = h_{33}$,
$ \left\langle (0,(g_{32}-h_{32}),h_{33},0,0,0,0,g_{38}),\, (1,0,0,-1,0,0,0,0) \right\rangle = 0$,
$ \left\langle (0,(g_{32}-h_{32}),h_{33},0,0,0,0,g_{38}),\, (1,0,0,0,-1,0,0,0) \right\rangle = 0$,
$ \left\langle (0,(g_{32}-h_{32}),h_{33},0,0,0,0,g_{38}),\, (1,0,0,0,0,-1,0,0) \right\rangle =0$,
$ \left\langle (0,(g_{32}-h_{32}),h_{33},0,0,0,0,g_{38}),\, (1,0,0,0,0,0,-1,0) \right\rangle = 0$,
$ \left\langle (0,(g_{32}-h_{32}),h_{33},0,0,0,0,g_{38}),\, (0,0,0,0,0,0,0,1) \right\rangle =g_{38} $.
\end{flushleft}

Now we calculate the limit to find the fix point associated to the face, $ \tau$ see formula in \cite{Brasselet}, thus,

\begin{equation}
X_{(0,(g_{32}-h_{32}),h_{33},0,0,0,0,g_{38})} =(\lim_{Z \longrightarrow 0}Z^{-(h_{33}+g_{38})},\lim_{Z \longrightarrow 0}Z^{-(g_{32}-h_{32}) }, \\ 

\lim_{Z \longrightarrow 0}Z^{h_{33}},  \lim_{Z \longrightarrow 0}Z^{0},\lim_{Z \longrightarrow 0}Z^{0},\lim_{Z \longrightarrow 0}Z^{0},
\lim_{Z \longrightarrow 0}Z^{0},\lim_{Z \longrightarrow 0}Z^{g_{38}}).
\end{equation}

\begin{equation}
X_{(0,(g_{32}-h_{32}),h_{33},0,0,0,0,g_{38})} =(0,0,0,1,1,1,1,0).
\end{equation}

This fix point $(0,0,0,1,1,1,1,0)  $ is true, if and only if the following constraints in the kinetic orders are also true,\\

\begin{equation}
h_{33}+g_{38} \,\leq \,0 ,\\
\end{equation}
\begin{equation}
g_{32}-h_{32} \,\leq  \,0  ,\\
\end{equation}
\begin{equation}
h_{33} \,\geq\, 0  ,\\
\end{equation}
\begin{equation}
g_{38}\,\geq\,0  .
\end{equation}

In the same manner, we calculate the fix points associated to the other faces or rays of cone, $(0,g_{32},0,0,0,0,0,g_{38})$ and $ (0,g_{32},h_{33},0,0,0,0,0) $, with fix points respectively given by, $(0,0,1,1,1,1,1,0)  $ with constraints, $ g_{38}=0 $ and $ g_{32} \,\leq\, 0 $, for $(0,g_{32},0,0,0,0,0,g_{38})  $ the fix point is $ (1,1,1,0,0,0,0,0) $ with constraints $ h_{33} =0$ and $ h_{32}\,\leq\,0 $.\\

We assume that all cones we work here with, contain the zero vector as a face, and always the fix point associated to this point is given by, $ X_{(0,...,0)}=(1,1,..,1) $. \\

\textbf{Explanation of fix points in gene and metabolic networks.}\\\\

An interpretation of the following formula $ X_{b_{i}} = \lim_{Z \longrightarrow 0} Z^{<b_{i},h_{j}>}  $ is as follows. For computation of fix points in gene networks, this limit represents when the biomass of metabolites and proteins in a genetic and metabolic network, the production or degradation of them, or an infinitesimal interpretation, we write, $ X_{1}\, \longrightarrow \,0,...,X_{8}\,\longrightarrow \,0$ for the variables in our example. In the case of the fix point $(0,0,0,1,1,1,1,0)  $, it means when the stimulus $X_{8}\,\longrightarrow \,0  $ disappears, also the messenger $ X_{1}\,\longrightarrow \,0  $ stops to synthesize biomass for the regulator, $X_{2}\,\longrightarrow \,0 $, and as a consequence, we also have $X_{3}\,\longrightarrow \,0  $. Then, if we see in Figure 1, the biomass in $ X_{7} $ stays at constant rate. These arguments are in agreement with the constraints calculated, previously. For example, if we take the equality of the constraints, it is equivalent to the analysis of the fix point, $(0,0,0,1,1,1,1,0)$. Hence,\\

\begin{equation}
 h_{33}+g_{38}= 0 ,\\
 \end{equation}
\begin{equation}
 g_{32}-h_{32} =0 ,\\
 \end{equation}
\begin{equation}
 h_{33} = 0,\\
 \end{equation}
\begin{equation}
 g_{38}= 0 .
\end{equation}

If $h_{33} = 0   $ and $ g_{38}= 0 $, the first equations are true. For the second equality, we have $g_{32}=h_{32}  $. If we substitute in the original equation $ \dot{X}_{3}= 0 $, in steady-state, it follows that,\\

\begin{equation}
\dot{X}_{3} = 0 = \alpha_{2,8} X^{g3,2}_{2} X^{0}_{8} - \beta_{3,2} X^{h3,2}_{2} X^{0}_{3} ,\\
\end{equation}
\begin{equation}
=\alpha_{2,8} X^{g3,2}_{2}- \beta_{3,2} X^{h3,2}_{2},\\
\end{equation}
\begin{equation}
\alpha_{2,8} X^{g3,2}_{2}= \beta_{3,2} X^{h3,2}_{2},\\
\end{equation}
\begin{equation}
\alpha_{2,8}= \beta_{3,2},\\
\end{equation}\\


\\
In the final equality, we obtain a steady-state condition for \( X_{3} \): the rate of production equals the rate of degradation, and the terms involving the stimulus \( X^{0}_{8} \) and the degradation of \( X^{0}_{3} \) vanish from the equation.

Based on the limit formula introduced above, we assume that it is possible to compute all the fixed points on the torus. For further details, see \cite{GEwald1993}, \cite{DCoxLittleOshea2004}. These computations are valid whenever the dual cone is \textbf{regular}.

The resulting fixed points are always of the form
\[
X_{b_{i}} = (0, 0, \ldots, 1, 1, 1, 0, 1, 0, \ldots, 1),
\]
where the entries with zero values correspond to species undergoing degradation or having null biomass, while the ones indicate species that retain biomass or are in production.

It is important to note that additional fixed points may exist on the torus, since the Hilbert basis is not unique. These alternative fixed points correspond to other possible states or configurations of the gene network, each associated with different elements of the basis.\\

%*\textbf{Linearized expression.}\\\\


\textbf{Linearized Expression}\\\\

It is important to emphasize that all the fixed points computed, denoted by \( X_{\tau} \), where \( \tau \) represents a face or ray of the cone, \textbf{do not depend on kinetic parameters}. Instead, they depend solely on the \textbf{topology of the network} and the \textbf{signs of the interactions} between species. 

This property allows us to analyze the dynamics of gene networks in \textbf{topological coordinates}, such as 
\[
u = (u_{1}, u_{2}, u_{3}, u_{4}, u_{5}, u_{6}, u_{7}, u_{8}) \in (\mathbb{C}^{*})^{8}.
\]

In what follows, we apply the \textbf{Jacobian matrix} technique to linearize and parametrize the equations governing the regulator \( \dot{X}_{3} \). To do so, we select a fixed point for linearization. For a complete dynamical description of the network, this process must be carried out for each fixed point on the torus. However, here we focus on a single fixed point,
\[
X_{(0,\,g_{32}-h_{32},\,h_{33},\,0,\,0,\,0,\,0,\,g_{38})} = (0, 0, 0, 1, 1, 1, 1, 0),
\]
and derive the linearized expression as follows:


\begin{equation}
\dot{X}_{3}=J\left(\dfrac{X_{1},...,X_{8}}{u_{1},...,u_{8}} \right)_{(0,0,0,1,1,1,1,0)} \bullet (u_{1},...,u_{8}) ^{T}.
\end{equation}

Where $ \bullet $, represents the scalar product of vectors and $ T $, transposition of vectors, thus we have,

\begin{equation}
\dot{X}_{3}=\left(0,g_{32}(\alpha_{28}-\beta_{32}),g_{38}\alpha_{28}-\beta_{33}h_{33},0,0,\alpha_{28}g_{16},0,0 \right)*(u_{1},...,u_{8}) ^{T}  ,\\\\\\\\
\end{equation}

\begin{equation}
\dot{X}_{3}=(g_{32}(\alpha_{28}-\beta_{32}))*u_{2}+(g_{38}\alpha_{28}-\beta_{33}h_{33})*u_{3}+\alpha_{28}g_{16}  *u_{6}.
\end{equation}

The benefits from this analysis is that we only need this analytical linear expression to know locally the behaviour of the inducer $ X_{3} $, we do not need to know the complete analytical solution of the set of S-systems of the network, as we can see in the last section.


\section{Topological Environment and Orthogonal Enzyme Kinetics}

In this section, we provide a formal definition of the \textbf{Topological Environment}. Before doing so, we first explain the underlying methodology using the gene circuit example developed in the previous section.

We model genetic and metabolic networks using S-systems, which are described by highly nonlinear differential equations. Solving the full set of equations analytically is computationally intensive and presents a significant challenge. The central philosophy of the Topological Environment approach is to focus on the \textbf{local neighborhood} of a given species, metabolite, or biomolecule within a gene network. Specifically, we aim to identify the interacting components and regulatory elements that directly surround a target species.

Instead of solving the entire system, we isolate and solve only a relevant \textbf{subset} of equations. If the complete system is desired, we can reconstruct it by \textbf{gluing together} all such local subsets. This method is analogous to techniques in algebraic geometry: each subset of information is represented as a \textbf{cone}, and the complete collection of these subsets is called a \textbf{fan}. In algebraic terminology, each piece is referred to as an \textbf{affine chart}, and the union of all charts forms an \textbf{algebraic variety} specifically, a \textbf{toric variety} in our context.

Within gene network modeling, each set of kinetic parameters associated with a biomolecule provides \textbf{local information}, which we associate with a cone or affine chart. These are linked to various biological events occurring in the network. We refer to such events as \textbf{environment circuits}, which represent regulatory mechanisms surrounding a given biomolecule. These environment circuits, interpreted through the lens of algebraic geometry, define the \textbf{Topological Environment} of the biomolecule and determine both its steady-state behavior and its dynamic response within the network.

Biomolecules participate in various biochemical reactions, and thus, gene and metabolic networks must be analyzed from multiple perspectives or \textbf{scenarios}. In our approach, we focus on a single biomolecule within the network, which is governed by a specific nonlinear differential equation (an S-system). This equation defines the particular \textbf{scene} or context for our analysis.

Temporarily setting aside the influence of other species, we examine the \textbf{local environment} of the target biomolecule. This local environment is represented mathematically by the \textbf{dual cone} associated with the support of the polynomial describing the S-system. Each row in the matrix corresponding to this dual cone encodes a \textbf{local event} or a distinct piece of regulatory or biochemical information within the network.

A schematic representation of this concept is provided in the following figures.


\begin{figure}[H]
\centering
\makebox{\includegraphics[height=0.55in]{Circuit2dpi.png}}
$ \oplus$
\makebox{\includegraphics[height=0.55in]{Circuit3dpi.png}}
$ \oplus $
\makebox{\includegraphics[height=0.55in]{Circuit4dpi.png}} 

\caption{\label{fig_simvarex} Topological environment and environment circuits associated to an Inducer \textbf{$X_{3}$}.}
\end{figure}

\textbf{Orthogonal Enzyme Kinetics from the Dual Cone} \\

As previously mentioned, the absence of known parameters in genetic–metabolic networks presents a major challenge for their analysis. The methodology of the \textbf{Topological Environment} offers an alternative approach to address this issue by studying the \textbf{exponent space} associated with S-system equations. In this technique, the focus is not initially on solving the system for the variables, but rather on analyzing the structure of the exponents variable resolution is deferred to a later stage.

We begin by analyzing the exponents in the topological environment, constructing a matrix representation from them, as outlined earlier.

The key advantage of analyzing this matrix associated with kinetic orders is that knowledge of the exact kinetic parameters is not required. In the topological environment, we work with the \textbf{dual cone} \( \sigma^{\vee} \) derived from the original cone \( \sigma \), and the corresponding matrix does not rely on kinetic parameters. This is a consequence by definition, lattice vectors belonging to the dual cone satisfy the algebraic condition:

\begin{center}
\(\left\langle b, \sigma \right\rangle \geq 0,\quad \text{for all } b \in \sigma^{\vee}.\)
\end{center}

One natural way to select a vector \( b \in \mathbb{Z}^{n} \) in the dual cone is to choose an \textbf{orthogonal vector}, i.e., a vector satisfying the stricter condition:

\begin{center}
\(\left\langle b, \sigma \right\rangle = 0.\)
\end{center}

For example, consider the lattice vector of kinetic orders associated with the inducer \( X_{3} \) from the previous section, which is given by:
\[
(0, g_{32}, 0, 0, 0, g_{36}, 0, g_{38}).
\]
Here, the kinetic orders \( g_{32} \) and \( g_{38} \) correspond to an enzyme-mediated reaction: \( g_{38} \) is the order associated with the substrate, while \( g_{32} \) is the order related to the catalyst \( X_{2} \), which facilitates the production of the inducer \( X_{3} \). The kinetic order \( g_{36} \) is associated with binding sites involved in the expression of the gene \( X_{6} \).

An orthogonal vector to this kinetic vector is, for instance:

\begin{center}
\(\left\langle (1, 0, 1, 1, 1, 1, 1, 0),\ (0, g_{32}, 0, 0, 0, g_{36}, 0, g_{38}) \right\rangle = 0.\)
\end{center}

This orthogonal vector, \( (1, 0, 1, 1, 1, 1, 1, 0) \sim (X_{1}, 0, X_{3}, X_{4}, X_{5}, X_{6}, X_{7}, 0) \), is independent of the kinetic orders themselves. Instead, it identifies the positions of the kinetic orders relative to neighboring reactions. In other words, one way to construct an orthogonal vector is by including all \textbf{complementary reactions} such as the production of transcripts \( X_{1} \) and \( X_{4} \), the expression of gene \( X_{6} \) via the inducer \( X_{3} \), and the generation of metabolites \( X_{5} \) and \( X_{7} \).

In this sense, every lattice vector \( b \in \mathbb{Z}^{n} \) in the dual cone that satisfies
\[
\left\langle b, \sigma \right\rangle = 0
\]
defines an instance of \textbf{orthogonal enzyme kinetics} with respect to the enzyme reactions encoded by the lattice vectors of the cone \( \sigma \).

As another example, consider the alternative orthogonal vector:
\[
(1, 0, 1, 0, 1, 1, 0, 0),
\]
which captures only the dynamics of gene expression for \( X_{6} \), including the production of transcript \( X_{1} \) via the inducer \( X_{3} \) and regulator \( X_{5} \). This constitutes another valid instance of orthogonal enzyme kinetics. A schematic illustration of this scenario is provided in the supplementary material.\\


The complete analysis of the \textbf{Topological Environment} in genetic–metabolic networks is fundamentally based on the family of \textbf{orthogonal vectors} discussed above. These vectors allow us to explore the system's structure without requiring prior knowledge of kinetic orders at least not until new differential expressions are introduced to describe the dynamics and steady-state behavior of the gene network.\\

These differential expressions can be organized into three main components:

\begin{enumerate}
    \item A \textbf{subset of equations} selected from the original, complete system of equations governing the network.
    \item A new set of \textbf{purely linear algebraic equations}, derived from toric geometry representations (e.g., via the Hilbert basis or dual cones).
    \item A system of classical \textbf{ordinary differential equations (ODEs)}, obtained through the linearization of S-systems using the Jacobian matrix.
\end{enumerate}

We now formally summarize the new concepts developed thus far in the form of two definitions, employing the terminology of \textbf{toric combinatorial geometry} and \textbf{S-systems}. For further technical background on these definitions, we refer the reader to the classical literature \cite{GEwald1993}, \cite{BSturmfels1996}, \cite{MASavageauVoit1987}.

\begin{definition}\label{def:env_cone} \textbf{Environment Cone.}  
Let \( f \) be an S-system representing the rate law for the concentration change of a given species in a genetic and metabolic network. Let \( \mathrm{supp}(f) \) denote the \textbf{support} of the polynomial \( f \), and let \( \sigma \subseteq \mathbb{Z}^{n} \) be the cone generated by this support. Then \( \mathrm{supp}(f) \subseteq \sigma \). 

We define the \textbf{environment cone} associated with \( f \) as the \textbf{dual cone} \( \sigma^{\vee} \). By construction, we also have:
\[
\mathrm{supp}(f) \subseteq \sigma \subseteq \sigma^{\vee}.
\]
\end{definition}

\begin{definition}\label{def:top_env} \textbf{Topological Environment.}  
Let \( f_1, \ldots, f_n \in \mathbb{C}[X, X^{-1}] \), where \( X = (X_1, \ldots, X_n) \in \mathbb{C}^n \), be a collection of S-systems representing the rate laws for biochemical concentration changes of molecular species in a genetic–metabolic network. For each polynomial \( f_i \), let \( \sigma_i^{\vee} \subseteq \mathbb{Z}^{n} \) be the dual cone associated with the support \( \mathrm{supp}(f_i) \).

We define the \textbf{Topological Environment} of the genetic–metabolic network as the \textbf{fan}:
\[
\Sigma := \bigcup_{i=1}^{n} \sigma_i^{\vee} \subseteq \mathbb{Z}^{n},
\]
where \( \Sigma \) captures the collection of all environment cones and encodes the local topological structure surrounding each species via the associated environmental variables \( X_i \in \mathbb{C} \).
\end{definition}

Within this fan i.e., the Topological Environment of the genetic and metabolic network, we compute \textbf{fixed points} to analyze and solve the dynamics of molecular species within the system.

%%%%%%%%%%%%%%%%%% Toric S-systems %%%%%%%%%%%%%%%%

In summarizing the previous results, we provide a formal statement in the following theorem. 
The relevance of this result is that it clarifies some conjectures in the context of the S-system approach. Firstly, we remember that the toric ideal, is a set of prime ideals or irreducible binomials. If $I:=< X^{a - X^{b}}>, a, b \in \mathbb{Z}^{n}$, and there exists a matrix $A$ such that $A(a-b)^{T}=$, then the lattice vector $(a-b)$ belongs to the kernel of $A$.\\

\begin{theorem}
    \textbf{(Toric S-system)}: 
Let 
\[
\dot{X}_{i}=  \alpha_{ip_{i}} \prod_{j=1}^{m} X_{j}^{g_{ip_{i}}} 
-  \beta_{iq_{i}} \prod_{j=1}^{m}X_{j}^{h_{iq_{i}}}, 
\quad i=1,\dots,n,
\]
be a set of S-system equations. 
Then, there exists a toric ideal, namely
\[
I_{H} = \langle X^{a} - X^{b} \; : \; H(a-b)^{T} = 0 \in \mathbb{Z}^{n}, , a, b \in \mathbb{Z}^{n} \ \rangle,
\]
where $H$ is a Hilbert basis with $\det(H)= \pm 1$. 
Moreover, the toric binomials $(X^{a} - X^{b})$ generate the S-system equations, that is, $\dot{X}_{i} \in I_{H}.$$
\end{theorem}
The proof of this fact is provided in the support information. This result is consequence from the classical result by Hironaka´s theorem, see\cite{MFAtiyah1970} and Hilbert basis, that S-systems equations can be emmbeded into toric ideals, where Hilbert basis $det(H)=\pm1$, reveals a resolution in dymanics for molecular networks, in the case when the $det(G-H)=0$.
%%%%%%%%%%%%%%%%%%%%%%%%%%%%%%%%%%%%%%%%%%%%%%%%%%%%%%%%%%%%%%%%%%%%%%%%%%%%%%
\subsection{Computing Dominant Fluxes in the Topological Environment}

Consider the following expression, obtained after parametrization within the Topological Environment. Let this parametrization be associated with a fan \( \Sigma \) or, more locally, with an environment cone \( \sigma^{\vee} \) corresponding to a toric variety \( X_{\sigma} \).\\


%\subsection{Computing dominant fluxes in Topological Environment}

%Given the following expression once parametrized in topological environment, for some fan $ \sum $ or environment cone $ \sigma^{\vee} $ associated to a toric variety $ X_{\sigma} $.\\
\begin{equation}
\dot{X}_{3}(u_{1},u_{2},u_{3},u_{4})=c_{1}*u_{6} + c_{2}*(u^{2}_{2}u^{2}_{3}u^{2}_{4}u^{2}_{6})u_{6}+c_{3}*u^{2}_{4}u_{6}-c_{4}*u_{3}u^{2}_{4}u_{6}.
\end{equation}\\
%\textbf{Theory of branches and Newton polytope, [].}\\
We define the \textbf{dominant branch} (or \textbf{main branch}) of the expression \( \dot{X}_{3}(U) \) as follows.

First, we compute the \textbf{support} of the parametrized expression \( \dot{X}_{3}(U) \), where the system has been transformed into toric coordinates. Then, for each element \( a^{*}_{i} \) in the \textbf{Hilbert basis} computed for the dual cone \( \sigma^{\vee} \), we evaluate all \textbf{projections} of the support vectors \( m_{j} \) onto \( a^{*}_{i} \). These projections are given by the scalar products: \[\left\langle a^{*}_{i},\, m_{j} \right\rangle,\]
where index \( i \) runs over the elements of the Hilbert basis and \( j \) over the elements of the support.

The dominant branch is identified by selecting the terms (or monomials) in the support that yield the \textbf{maximum projection values} for a fixed \( a^{*}_{i} \). These projections reflect which reactions or regulatory terms dominate the local dynamics of \( \dot{X}_{3}(U) \) in the Topological Environment.


\begin{flushleft}
$ Supp(\dot{X}_{3}(U))=\left\lbrace  m_{1}=(0,0,0,0,0,1), m_{2}=(0,2,2,2,0,3), m_{3}=(0,0,0,2,0,1), m_{4}=(0,0,1,2,0,1) \right\rbrace . $
\end{flushleft}

We compute all the \textbf{projections} associated with these faces by evaluating the scalar products between the rays \( m_1, m_2, m_3, \) and \( m_4 \), and the elements of the Hilbert basis \( a^{*}_{j} \). That is, we compute all scalar products of the form:
\[
\left\langle a^{*}_{j},\, m_{i} \right\rangle,
\]
where \( j \) indexes the elements of the Hilbert basis and \( i = 1, 2, 3, 4 \) corresponds to the rays \( m_{i} \) in the support of the expression \( \dot{X}_{3}(U) \).

%\begin{comment}
\begin{flushleft}
$ \left\langle a^{*}_{1}, m_{1} \right\rangle=0  $, and $ \left\langle a^{*}_{1}, m_{2} \right\rangle=0 $,\\
$ \left\langle a^{*}_{2}, m_{1} \right\rangle=0 $, and $ \left\langle a^{*}_{2}, m_{2} \right\rangle=2 $\\
$ \left\langle a^{*}_{3}, m_{1} \right\rangle=0 $, and $ \left\langle a^{*}_{3}, m_{2} \right\rangle=4 $\\
$ \left\langle a^{*}_{4}, m_{1} \right\rangle=0 $, and $ \left\langle a^{*}_{4}, m_{2} \right\rangle=8 $.\\
$ \left\langle a^{*}_{5}, m_{1} \right\rangle=0 $, and $ \left\langle a^{*}_{5}, m_{2} \right\rangle=0 $.\\
$ \left\langle a^{*}_{6}, m_{1} \right\rangle=-1 $, and $ \left\langle a^{*}_{6}, m_{2} \right\rangle=1 $.\\
\end{flushleft}
For the vectors, $m_{3}$ and $ m_{4} $, we get the minimal projections.\\

\begin{flushleft}
%\begin{center}
$ \left\langle a^{*}_{1}, m_{3} \right\rangle=0  $, and $ \left\langle a^{*}_{1}, m_{4} \right\rangle=0 $,\\
$ \left\langle a^{*}_{2}, m_{3} \right\rangle=0 $, and $ \left\langle a^{*}_{2}, m_{4} \right\rangle=0 $\\
$ \left\langle a^{*}_{3}, m_{3} \right\rangle=0 $, and $ \left\langle a^{*}_{3}, m_{4} \right\rangle=1 $\\
$ \left\langle a^{*}_{4}, m_{3} \right\rangle=2 $, and $ \left\langle a^{*}_{4}, m_{4} \right\rangle=4 $.\\
$ \left\langle a^{*}_{5}, m_{3} \right\rangle=0 $, and $ \left\langle a^{*}_{5}, m_{4} \right\rangle=0 $.\\
$ \left\langle a^{*}_{6}, m_{3} \right\rangle=-1 $, and $ \left\langle a^{*}_{6}, m_{4} \right\rangle=0 $.\\
%\end{center}
\end{flushleft}
%\end{comment}
%%%%%%%%%%%%%%%%%%%%%%%%%%%%%%%%%%%%%%
The \textbf{minimal branch} \( \gamma \) that intersects the faces of the cone is defined by the following scalar projections:
\[
\left\langle a^{*}_{6},\, m_{3} \right\rangle = -1 \quad \text{and} \quad \left\langle a^{*}_{6},\, m_{1} \right\rangle = -1.
\]
Therefore, the only terms that contribute to the analysis of the equation \( \dot{X}_{3}(U) \) along this branch are those corresponding to the rays \( m_{1} \) and \( m_{3} \).

For a detailed study of \textbf{bifurcations in dynamical systems}, and the role of branches in \textbf{Newton polytopes} and \textbf{cones with toric resolution}, we refer the reader to the classical works \cite{TFukui2000}, \cite{GEwald1993}, \cite{BSturmfels1996}.

Thus, we obtain:\\

%The minimal branch $ \gamma $ which intersects the faces is defined by the projections, with $\left\langle a^{*}_{6}, m_{3} \right\rangle=-1  $ and $ \left\langle a^{*}_{6}, m_{1} \right\rangle=-1 $, therefore the only terms to study the equation $ \dot{X}_{3}(U) $ are in $ m_{1} $ and $ m_{3} $. For a deep study of  bifurcation in dynamical systems and branches in Newton polytopes and cones with toric resolution, see, \cite{TFukui2000}, \cite{GEwald1993}, \cite{BSturmfels1996}, thus, we have.\\

\begin{equation}
\dot{X}_{3}(U)_{\gamma}=c_{1}*u_{6} + c_{3}*u^{2}_{4}u_{6} .
\end{equation}

%As we can see, to analyze this expression, is equivalent to analyse the dominant fluxes in S-systems, using the tools in bifurcation theory and toric algebraic geometry, and we use this expression in the last section for the reproduction of oscillatory phenotypes, \cite{LomnitzSavageau2014}.

As we can see, analyzing this expression is equivalent to studying the \textbf{dominant fluxes} in S-systems, using tools from \textbf{bifurcation theory} and \textbf{toric algebraic geometry}. This formulation will be employed in the final section to reproduce \textbf{oscillatory phenotypes}, as discussed in \cite{MASavageau2001}.


\section{Results: Dynamics in Topological Environment}

\textbf{Linearization of Nonlinear Dynamical Systems via Fixed Points on algebraic trous} \\

In this example, we describe a gene network consisting of two genes and two regulatory elements, figures 3, 4 and 5. The network operates under multiple transcriptional control modes that is, gene expression is governed by a variety of mechanisms including repression, activation, and their combinations. The mechanism of gene regulation is governed by the integers $\delta_{i}$ and $\pi_{i}$ described ahead.

The primary goal of this section is to derive an \textbf{analytical expression} for the variable \( X_{4} \) in \textbf{logarithmic space}, and to reproduce simulations of the \textbf{oscillatory quantitative phenotypes} associated with these gene circuits. This system was previously studied by M.A. Savageau in \cite{LomnitzSavageau2014}. These mechanisms of gene regulation are involved in a variety of phisiologies in natural circadian riythms in Human and bacteria, \cite{Benner}, \cite{Kim} and \cite{Tigges}.

\vspace{1em}
\noindent
\textbf{Example: Reproduction of Oscillatory Quantitative Phenotypes.}\\

\begin{figure}[htbp]
\centering
\makebox{\includegraphics[height=2.2in]{Circuit5dpi.png}}
\caption{\label{fig_simvarex} 
%Schematic representation of oscillatory gene circuit with six elements. $X_{4}$ antagonizes the activity for $X_{1}$ and $X_{3}$ also is repressed by protein $X_{2}$ (Case 1).
Schematic representation of an oscillatory gene circuit with six elements. In this configuration (Case 1), \( X_{4} \) antagonizes the activity of \( X_{1} \), and \( X_{3} \) is also repressed by the protein \( X_{2} \).
}
\end{figure}

\begin{figure}[htbp]
\centering
\makebox{\includegraphics[height=2.2in]{Circuit6dpi.png}}
\caption{\label{fig_simvarex}
Schematic representation of an oscillatory gene circuit with one control mode. In this configuration, \( X_{2} \) primarily activates \( X_{1} \), after \( X_{4} \) represses the activity of \( X_{2} \) (Case 2). In a second control mode, \( X_{4} \) primarily represses the activity of \( X_{1} \), and is subsequently antagonized by the protein \( X_{2} \) (Case 3).}
\end{figure}

\begin{figure}[htbp]
\centering
\makebox{\includegraphics[height=2.2in]{Circuit7dpi.png}}
\caption{\label{fig_simvarex}
Schematic representation of an oscillatory gene circuit with two control modes. In this diagram, only one case of the two-control-mode configurations is illustrated (Case 4), where \( X_{2} \) activates \( X_{1} \), and \( X_{4} \) represses \( X_{2} \). Additionally, \( X_{3} \) is activated by \( X_{2} \), while \( X_{4} \) antagonizes the activity of \( X_{2} \), as shown in the figure. Three additional control mode cases can be represented within this circuit: \textbf{Case 5:} The repressor \( X_{4} \) primarily antagonizes \( X_{1} \) and is subsequently repressed by both \( X_{2} \) and \( X_{3} \). In this case, \( X_{3} \) is regulated in the same manner as in Case 4. \textbf{Case 6:} \( X_{2} \) activates \( X_{1} \), \( X_{4} \) represses \( X_{2} \), and then \( X_{4} \) also represses \( X_{3} \) after being repressed by \( X_{2} \). \textbf{Case 7:} The protein \( X_{4} \) directly represses \( X_{1} \) after being repressed by \( X_{2} \). The control mode for \( X_{3} \) remains the same as in Case 6.}
\end{figure}

In Savageau and Lomnitz \cite{LomnitzSavageau2015}, the following variables are considered: $X_{2i}=x_{2i}/(K^{0}_{2i}), \: i=1,2.$, and\\
\begin{equation}
%\begin{center}
X_{2i-1}= \left(\dfrac{(\beta^{0}_{2i-1})(\sqrt{\rho^{0}_{2i-1}})}{\alpha^{0}_{2i-1}} \right)x_{2i-1}, \, i=1,2., 
%\end{center}
\end{equation}
The concentrations of the immature activator and repressor, denoted \( X_{A} \) and \( X_{R} \), are normalized with respect to their steady-state values. In other words, the regulator concentrations are scaled by their respective \textbf{DNA dissociation constants}. 

In the vicinity of their promoter regions, it is assumed that both promoters contain identical pairs of binding sites for the repressor–DNA and activator–DNA complexes. That is,
\[
K_{2R} = K^{0}_{2} \quad \text{and} \quad K_{4A} = K^{0}_{4}.
\]
The parameters used in the model are explained in detail below.\\\\

We begin the analysis of this gene circuit using the methodology of \textbf{Topological Environment Points}, selecting only a \textbf{subset of S-system equations} from the original system for analysis of our methodology. The complete set of S-system equations for this example is provided in \cite{LomnitzSavageau2014}.\\

\begin{equation}
\dot{X}_{3}=\left[  \alpha_{3} (\beta^{3}_{0} + \mu^{0}) \sqrt{\rho^{0}_{3}} (\alpha_{3}^{0})^{-1} \right] \lbrace (\rho_{3})^{- \pi_{3}}(X_{6})^{-1} +
(K_{2R})^{-g_{32}}(K_{2}^{0})^{g_{32}} (X_{2})^{g_{32}}(X_{6})^{-1} +\\

\delta_{3}(\rho_{3})^{-1} (K_{4})^{-g_{34}}(K_{4}^{0})^{g_{34}}X_{4}^{g_{34}} (X_{6})^{-1}  \rbrace - (\beta_{3}+ \mu) X_{3} ,\\\\
\end{equation}

\begin{equation}
\dot{X}_{R}= (\beta^{0}_{R} + \mu^{0})\left[ \dfrac{\alpha_{R}}{\alpha^{0}_{R}} \right]  X_{3} - (\beta_{R} + \mu) X_{R},\\\\
\end{equation}

\begin{equation}
\dot{X}_{4} = (\beta^{0}_{4} + \mu^{0}) \left( \dfrac{\beta_{R}}{\beta^{0}_{R}}\dfrac{\alpha_{3}^{0} n_{4}^{0}}{(\beta_{3}^{0} + \mu^{0})\sqrt{\rho_{3}^{0}} K_{4}^{0}} \right) X_{R} - (\beta_{4} + \mu)X_{4}.\\\\
\end{equation}

We now introduce two additional expressions, related to the behavior of the regulator and the activator. These expressions are derived from \textbf{Michaelis–Menten type relationships} that describe enzyme kinetics in the gene circuit under consideration. For further details, see Savageau's work in \cite{LomnitzSavageau2014}.
They are the kinetic parameters $ K_{2R} $ and $ K_{4A}$ are defined as:\\\\\\

\begin{equation}
K_{4A}= \dfrac{\left[  \dfrac{\alpha_{3}^{0} n_{4}^{0}}{(\beta_{3}^{0} + \mu^{0})\sqrt{\rho_{3}^{0}}} \right]}{\left\lbrace   \left( \delta_{1} \left[ \dfrac{\alpha_{1}^{0} n_{2}^{0}}{(\beta_{1}^{0} + \mu^{0})\sqrt{\rho_{1}^{0}} K_{2}^{0}} \right]^{g_{12}}  + 1 - \pi_{1}  \right) \sqrt{\rho_{1}^{0}} - \pi_{1} \right\rbrace^{\dfrac{1}{g_{14}}}};
\end{equation}\\\\

\begin{equation}
K_{2R}=\dfrac{   \left[  \dfrac{ \alpha_{1}^{0} n_{2}^{0}  }{(\beta_{1}^{0}  + \mu^{0}) \sqrt{\rho_{1}^{0}} } \right] \sqrt{\rho_{3}^{0}}^{\dfrac{1}{g_{32}}}}{ \left\lbrace   \delta_{3} \left[  \dfrac{\alpha_{3}^{0} n_{4}^{0}}{(\beta_{3}^{0}  + \mu^{0})\sqrt{\rho_{3}^{0}} K_{4}^{0} }  \right]^{g_{34}}   + (\pi_{3} - 1)\sqrt{\rho_{3}^{0}} + \pi_{3} \right\rbrace^{\dfrac{1}{g_{32}}}   } . 
\end{equation}\\\\\\

These kinetic parameters depend on binary regulatory parameters, denoted \( \pi_{i} \) and \( \delta_{i} \), which govern the gene regulation mechanisms in the gene circuits in the figures. These parameters, together with the additional constants used in the above equations, will be explained in detail below. For a comprehensive derivation and interpretation of these parameters, we refer the reader to the original work by Savageau \cite{LomnitzSavageau2014}. In this study, we make use of the equations already deduced and provide a brief explanation of their structure and role within the system.\\

For this system, the \textbf{condition lattice vector} is defined as
\[
g = (g_{12},\, g_{14},\, g_{32},\, g_{34}) \in \mathbb{Z}^{4},
\]
where each \( g_{ij} \) represents a \textbf{kinetic parameter} associated with the \textbf{cooperativity} of either repression or activation. These parameters correspond to the number of \textbf{binding sites} occupied by the respective regulatory proteins at the promoter regions of the genes.\\

The constants \( \alpha_{i}, \beta_{i}, \mu^{0}, \rho_{i}^{0}, n_{i}^{0} \in \mathbb{R} \) represent the following biological quantities:  
\begin{itemize}
  \item \( \alpha_{i} \): maximum rates of transcription,  
  \item \( \beta_{i} \): rate constants for mRNA and tagged protein degradation at mid-level expression,  
  \item \( \mu^{0} \): dilution rate due to exponential cellular growth,  
  \item \( \rho_{i}^{0} \): regulatory capacities,  
  \item \( n_{i}^{0} \): ratio of protein to mRNA concentrations.
\end{itemize}

The constants \( K_{2}, K_{4}, K_{2R}, K_{4A} \in \mathbb{R} \) denote concentrations of regulatory proteins required for \textbf{half-maximal binding} to control sequences in the DNA:
\begin{itemize}
  \item \( K_{2} \) and \( K_{4} \): binding near their respective promoter regions,  
  \item \( K_{2R} \) and \( K_{4A} \): binding near the promoter regions of other genes.
\end{itemize}

For further details on the definitions and derivations of these parameters, see the original work by J. Lomnitz and M. Savageau \cite{LomnitzSavageau2014}.\\

The parameters \( \pi_{i}, \delta_{i} \in \mathbb{Z}_{2} \), for \( i = 1, 2, 3, 4 \), are binary variables that define the different \textbf{modes of regulatory control} for the activator and repressor proteins in the gene and metabolic networks, see the figures 3, 4 and 5.

\begin{itemize}
    \item When \( \pi_{i} = 1 \), the system behaves as an \textbf{activator-primary} system: the default state of the promoter is inactive, and activation is required to initiate gene expression. The activator serves as the primary regulatory input to enhance transcription.
    
    \item When \( \pi_{i} = 0 \), the system behaves as a \textbf{repressor-primary} system: the promoter is fully active in the absence of regulatory inputs, and repression is required to reduce gene expression from its default active state.

    \item The parameter \( \delta_{i} = 1 \) indicates that the gene is regulated by \textbf{two regulators} (a double-regulator network), while \( \delta_{i} = 0 \) corresponds to regulation by \textbf{a single regulator}.
\end{itemize}

The solutions of the S-system described previously, under various combinations of control modes defined by \( \pi_{i} \) and \( \delta_{i} \), are illustrated in the figures, 3, 4 and 5.

\vspace{1em}
We now apply the principles of the \textbf{Topological Environment} methodology. One key aspect of this framework is that it allows for a \textbf{localized analysis} of the S-system equations.

In this gene network, our primary interest lies in describing the dynamics of the \textbf{repressor-regulator} \( X_{4} \), and in demonstrating the existence of an \textbf{oscillatory quantitative phenotype} or \textbf{sustained oscillatory region}.

To achieve this, we focus exclusively on the equation for \( \dot{X}_{3} \), since this variable is directly linked to the immature repressor \( X_{R} \), and thereby to the regulator \( X_{4} \). We perform a topological environment analysis of this equation, with the goal of obtaining an analytical expression for the dynamics of the regulator \( X_{4} \).


The cones \( \sigma_{\dot{X}_{3}} \) and the associated \textbf{environment cone} \( \sigma^{\vee}_{\dot{X}_{3}} \) are constructed following the methodology of the Topological Environment framework.\\\\
\vspace{1em}
\noindent
\textbf{Support Associated with \( \dot{X}_{3} \) in Matrix Representation}


%The cones $ \sigma_{\dot{X}_{3}} $ and environment cone $\sigma^{\vee}_{\dot{X}_{3}}   $, associated are given below, following the methodology,\\\\

%\textbf{Support associated to $ \dot{X}_{3}$ in a matrix representation}.\\

$Supp(\dot{X}_{3})=      
\begin{pmatrix}

0&2&0&0&0&-1\\

0&0&1&0&0&0\\

0&0&0&2&0&-1\\

0&0&0&0&0&-1\end{pmatrix}  
$, \\\\

\textbf{Cone associated to $ \dot{X}_{3} $}.\\\\

$\sigma_{\dot{X}_{3}}=      
\begin{pmatrix}

0&2&0&0&0&-1\\

0&2&-1&0&0&-1\\

0&0&1&-2&0&1\\

0&0&0&2&0&0\\

0&0&0&0&0&-1
\end{pmatrix} 
  
$.\\

\textbf{Environment cone associated to $ \dot{X}_{3} $}.\\\\

$\sigma^{\vee}_{\dot{X}_{3}}=      
\begin{pmatrix}

0&0&1&0&0&-1\\

0&1&0&0&0&0\\

0&1&2&0&0&0\\

0&1&2&1&0&0\\

\end{pmatrix}  
$.\\\\

\begin{figure}[htbp]
\centering
\makebox{\includegraphics[height=1.1in]{Circuit8dpi.png}}
\caption{\label{fig_simvarex} Topological environment associated to the face $ (0,1,2,1,0,0) $.}
\end{figure}

\begin{figure}[htbp]
\centering
\makebox{\includegraphics[height=1.1in]{Circuit9dpi.png}}
\caption{\label{fig_simvarex} Topological environment associated to the face $ (0,1,2,0,0,0) $.}
\end{figure}

\begin{figure}[htbp]
\centering
\makebox{\includegraphics[height=1.1in]{Circuit10dpi.png}}
\caption{\label{fig_simvarex} Topological environment associated to the face $ (0,0,1,0,0,-1) $.}
\end{figure}

%\begin{figure}[htbp]
%\centering
%\makebox{\includegraphics[height=0.60in]{oscillatory34.png}}
%\caption{\label{fig_simvarex} Topological environment associated to the face $ (0,1,0,0,0,0) $.}
%\end{figure}

Hilbert Basis in its matrix representation for $ \dot{X}_{3} $, we get,\\\\

%\begin{center}
$ H_{\dot{X}_{3}}=      
\begin{pmatrix}

0&1&2&0&0&0\\
0&1&2&1&0&0\\
0&0&1&0&0&-1\\
0&1&0&0&0&0\\
0&1&1&0&0&0\\
1&0&0&0&0&0\\
0&0&0&0&1&0\\
-1&0&0&0&0&0\\
0&0&0&0&-1&0\end{pmatrix}  $\\\\

%\end{center}

We chose one sub-modular matrix $[H_{\dot{X}_{3}}] $, thus,\\\\

%\begin{center}
$ [H_{\dot{X}_{3}}]=      
\begin{pmatrix}

1&0&0&0&0&0\\
0&1&0&0&0&0\\
0&1&1&0&0&0\\
0&1&2&1&0&0\\
0&0&0&0&1&0\\
0&0&1&0&0&-1
\end{pmatrix}  $\\\\
%\end{center}

We can check the property $ det([H_{\dot{X}_{3}}])=-1 $. Now we take the transpose of this sub-matrix, with the purpose of finding a monomial parametrization, as follows,\\\\

%\begin{center}
$ [H_{\dot{X}_{3}}]^{T}=      
\begin{pmatrix}

1&0&0&0&0&0\\
0&1&1&1&0&0\\
0&0&1&2&0&1\\
0&0&0&1&0&0\\
0&0&0&0&1&0\\
0&0&0&0&0&-1
\end{pmatrix}  $\\\\
%\end{center}

Hence, we can write the monomial transformation of changing coordinates, using our usual map $ a_{i}\,\longrightarrow \, X^{a_{i}} $ (section 1), as follows,\\

\begin{equation}
 X_{1}= u_{1} ,  X_{2}= u_{2} u_{3}u_{4},  X_{3}= u_{3}u^{2}_{4}u_{6} ,
 X_{4}= u_{4} ,  X_{5}= u_{5} ,  X_{6}= u^{-1}_{6} .
\end{equation}

In this local toric affine chart (monomial coordinates) we parametrise the expression for the equation $\dot{X}_{3}$, where their coefficients are denoted with the constants, $ c_{1},\,c_{2},c_{3},\,c_{4} $, the parametrised expression, gives,\\

\begin{equation}
\dot{X}_{3}(u_{1},u_{2},u_{3},u_{4})=c_{1}*u_{6} + c_{2}*(u^{2}_{2}u^{2}_{3}u^{2}_{4}u^{2}_{6})u_{6}+c_{3}*u^{2}_{4}u_{6}-c_{4}*u_{3}u^{2}_{4}u_{6}.
\end{equation}

Now we only consider the expression for the dominant flux $\dot{X}_{3}(U)_{\gamma}  $, hence,\\

\begin{equation}
\dot{X}_{3}(U)_{\gamma}=c_{1}*u_{6} + c_{3}*u^{2}_{4}u_{6}.
\end{equation}

The fix points associated to the faces or rays into the cone $ \sigma $ in this circuit, are calculated in the same manner as in section 2., and we only write the results, $X_{(0,2,0,0,0,-1)} = (1,0,0,0,1,0)$,  $X_{(0,0,1,-2,0,1)} = (1,1,0,1,1,1)  $, $X_{(0,0,0,2,0,0)} = (1,1,1,0,1,1)  $, $X_{(0,0,0,0,-1)} = (1,1,1,1,1,0)  $, \\

We linearize the expression at the fixed point 
\[
X_{(0,0,1,-2,0,1)} = (1, 1, 0, 1, 1, 1).
\]
The rationale for this choice is closely tied to the role of the \textbf{activator} \( X_{4} \), and consequently, to the regulation of the \textbf{inducer} \( X_{3} \). This interaction is explicitly observed in the monomial term \( X_{3} X_{4}^{-2} \), which defines the \textbf{dominant flux} of interest in this system. Furthermore, at this fixed point, the coordinate corresponding to the variable \( X_{3} \) is zero. As discussed in Section~2, this indicates that the system is near a \textbf{regulatory transition} in which the degradation of the inducer is dominant, i.e.,
\[
X_{3} \longrightarrow 0.
\]
This behavior suggests that the system is close to a gene regulatory event where the expression of \( X_{3} \) is being significantly downregulated or fully suppressed.\\


Therefore, we get,

\begin{equation}
\dot{X}_{3}(U)_{\gamma}=\left(0,0,0,2*c_{3}, 0,(c_{1}+ c_{3})  \right)*(u_{1},...,u_{6})^{T} .\\
\end{equation}

\begin{equation}
\dot{X}_{3}(U)_{\gamma}=(2c_{3})*u_{4}+(c_{1}+c_{3})*u_{6}.
\end{equation}
On the other hand, we have the change of variables given by:\[X_{3} = u_{3} u_{4}^{2} u_{6}.\]
We compute the derivative of this expression using the product rule. Then, by comparing the result with the previously obtained linearized expression for \( \dot{X}_{3} \), we derive the corresponding system of \textbf{ordinary differential equations} (ODEs) governing the dynamics of the transformed variables. The resulting system is as follows:


\begin{equation}
u^{2}_{4}\dot{u_{6}} =0,\: 2u_{3}u_{6}\dot{u_{4}}= 2c_{3},\:
u^{2}_{4}\dot{u_{3}}= (c_{1}+c_{3}).
\end{equation}

We analyze the resulting system of \textbf{ordinary differential equations} for various values of the constants \( c_{1} \) and \( c_{3} \), in accordance with the oscillatory gene circuit design studied by Savageau \cite{LomnitzSavageau2014}. In our simulations, the condition \( c_{3} \neq 0 \) corresponds to the case \( \delta_{3} = 1 \), representing a \textbf{double-regulator network}, while \( c_{3} = 0 \) corresponds to \( \delta_{3} = 0 \), indicating a \textbf{single-regulator network}. Each of these scenarios is further combined with all possible values of the binary control mode parameter \( \pi_{i} \). It is important to emphasize that, under this methodology, the original nonlinear system is successfully represented by a transformed set of ordinary differential equations such as the one derived above enabling analytical and numerical exploration of the system’s dynamic behavior.

Finally, substituting in $X_{3}= u_{3}u^{2}_{4}u_{6}$, the differential equations above, are solved for two different cases, with $c_{3}=0 $ and $c_{3}\neq 0 $, to get two solutions for $X_{3}(t)  $ respectively.\\

For $\delta_{3}=0  $.

\begin{equation}
X_{3}(t)=c_{3}(c_{1}+c_{3})[(t_{0}-t) +u_{40}].
\end{equation}

For $\delta_{3}=1 $.

\begin{equation}
X_{3}(t)=c_{1}u_{60}[(t- t_{0}) +u^{2}_{40}u_{30}/c_{1}].
\end{equation}
%We use the analytical solution $ X_{3}(t)  $, substituting in $ \.{X}_{R} $.\\
As we can see in the last equations we can substitute the function $ X_{3}(t) $, for each case in $ \delta_{3} $, to solve for $ X_{R} $ in the equation \textbf{4.3}, and then we can solve for $ X_{4} $ in the corresponding ordinary differential equation.\\

We substitute into the differential equations the new variables or environment variables $ X^{*}_{R}, X^{*}_{4} $, in functions on topological environment, see supplements to check details.\\

The solution for the first equation in $X^{*}_{R}  $ is solved in Wolfram Alpha on-line, https://www.wolframalpha.com/ , for both cases. Hance,\\

\textbf{Solution for $X_{R}^{*}(t)$ with $\delta_{3}=0  $ and $\delta_{3}=1  $, respectively}.\\

\begin{equation}
 X_{R}^{*}(t)=\left[\pm  \dfrac{ib}{a^{2}} \mp\dfrac{bc}{a} \pm \dfrac{b*t}{a} + K_{1}*\exp(-ia*t) \right] .
\end{equation}


We define $ f=(\beta^{0}_{4} + \mu^{0}) \left( \dfrac{\beta_{R}}{\beta^{0}_{R}}\dfrac{\alpha_{3}^{0} n_{4}^{0}}{(\beta_{3}^{0} + \mu^{0})\sqrt{\rho_{3}^{0}} K_{4}^{0}} \right) $, in \textbf{4.4} and we solve $ X^{*}_{4} $.\\

We plot the Real parts and Imaginary parts of $X^{*}_{4}  $ for the seven cases associated to the gene circuits in the figures 3, 4 and 5, thus we reproduce the quantitative phenotypes given in, ref. \cite{LomnitzSavageau2014}, as shown in the plots below;\\\\

\textbf{Case 1.} Solution with $ f $ depends on $ K^{0}_{4} $ for a real value (we only plot the Imaginary part). \\

\begin{equation}
Im(X^{*}_{4}(t))= \left( \dfrac{bf*t}{ap} -\dfrac{bcf}{ap}- \dfrac{fK_{1}}{(a-p)}*\cos(a*t)-\alpha_{1}\sin(p*t) \right) .
\end{equation}

\textbf{Case 2.} Solution with $ f $ depends on $ K^{0}_{4} $ for a real value (we only plot the Real part).\\

\begin{equation}
Re(X^{*}_{4}(t))=\left( \dfrac{bf}{a^{2}p} + \dfrac{bf}{ap^{2}} + \dfrac{fK_{1}}{(a-p)}*\sin(a*t)+ \alpha_{1}\cos(p*t)\right).
\end{equation}

\textbf{Case 3.} Solution with $ f $ depends on $ K^{0}_{4} $ for a complex value, ( we only plot the Real part).\\

\begin{equation}
Re(X^{*}_{4}(t))=\left(- \dfrac{bf}{a^{2}p}-\dfrac{bf}{ap^{2}}-\dfrac{fK_{1}}{(a-p)}*\sin(a*t)+ \alpha_{1}\cos(p*t)\right).
\end{equation}

\textbf{Cases, 4 and 6}. Solution with two different values for $ f $ respectively and $ K^{0}_{4} $ a real value, (we only plot the Imaginary part) .\\

\begin{equation}
Im(X^{*}_{4}(t))=\left(- \dfrac{bf}{a^{2}p}-\dfrac{bf}{ap^{2}} +\dfrac{fK_{1}}{(a-p)}*\sin(a*t)- \alpha_{1}\sin(p*t)\right).
\end{equation}


In the same manner the solution is obtained in Mathematica Alpha on-line, as follows,\\

\textbf{Cases, 5 and 7}. Solution with two different values for $ f $ respectively and $ K^{0}_{4} $ a complex value, (we only plot the Real part) .\\
\begin{equation}
Re(X^{*}_{4}(t))=\left( \dfrac{bf}{a^{2}p} + \dfrac{bf}{ap^{2}}-\dfrac{fK_{1}}{(a-p)}*\sin(a*t)+\alpha_{1}\cos(p*t) \right) .
\end{equation}
%%%%%%%%
The simulations of all those equations are shown below, and they reproduce the \textbf{oscillatory quantitative phenotypes} of the gene circuit of the figures 3-5 and their mechanisms of gene expression control, in agreement with the values $ \pi_{i} $ and $  \delta_{i} $, we plot these functions for different free parameters and constants, which produces different magnitudes in the logarithmic functions.\\

%%%%%%%%%%%%%%%%%%%%%%%%%%%%%%%%%%%%%%%%%%%%%%%%%%%%%%%%%%%%%%%%%%%%%%%%%%%%%

\begin{figure}[h]
\centering
\makebox{\includegraphics[height=2.65in]{Simulations.png}}
\caption{\label{fig_simvarex} Sustained oscillatory phisiologies with kinetic paramters for: Case 1) $K_{4A3}=K_{4A2}/\sqrt{\rho_{01}}$, $K_{4AR}=\sqrt{-K_{4A3}}$. Case 2) Parameters $K_{4A3}=1, K_{4AR}=1$. Case 3) $K_{4A3}=K_{4A2}/\sqrt{\rho_{01}}*K_{4A1}*K_{4A1} +
\sqrt{\rho_{01}}$, $K_{4AR}=\sqrt{-K_{4A3}}$. Case 4) $K_{4A3}=1,
K_{4AR}=1$. Case 5)
$K_{4A3}=K_{4A2}/\sqrt{\rho_{01}}*K_{4A1}*K_{4A1} + \sqrt{\rho_{01}}$, $K_{4AR}=\sqrt{-K_{4A3}}$. Case 6) $K_{4A3}=1,
K_{4AR}=1$. Case 7) $K_{4A3}=K_{4A2}/\sqrt{\rho_{01}}*K_{4A1}*K_{4A1} + \sqrt{\rho_{01}}$, $K_{4AR}=\sqrt{-K_{4A3}}$. Molar concentration for $X_{4}$ regulator in logarithmic space is quantified in y-axis and time scale (hrs) in x-axis.}
\end{figure}
%%%%%%%%%%%%%%%%%%%%%%%%%%%%%%%%%%%%%%%%%%%%%%%%%%%%%%%%%
%\begin{figure}[H]
%\centering
%\makebox{\includegraphics[height=0.90in]{top_env5dpi.png}}
%\caption{\label{fig_simvarex} The allosteric system of end-product inhibition for a pathway of length $n=3$ serves as a model for biosynthetic pathways (see \cite{SavageauOptimal}), such as the biosynthesis of homoserine regulated by methionine (see EcoCyc pathways). The topological environment of this system is divided into three local charts, corresponding to three Hilbert bases.
%}
%\end{figure}

\subsection{A Toy example.}

Finally apply this methodology to the case of an allosteric system by end-product inhibition for a pathway of length $n=3$, which exhibits oscillatory behaviour. This system models a biosynthetic pathway, such as the biosynthesis of homoserine regulated by methionine (see EcoCyc pathways). In the following, we provide conditions in the topological environment for the nonlinear expression of $X_{1}$. Its local dynamics confirm this quantitative phenotype, (see Fig.~1). Here, the allosteric protein $X_{1}$ catalyzes the enzyme $X_{2}$, after which the enzyme $X_{3}$ antagonizes the activity of the allosteric protein through eight regulatory subunits ($g=8$) in its quaternary structure (orange points), as shown in \cite{SavageauOptimal}. A detailed version of this example is provided in the Supporting Information.


\section{On the Computational Complexity}

The issue of computational complexity is addressed in \cite{Henk}. 
In general, the computation of a Hilbert basis is NP-hard. 
However, in cases where the Hilbert basis is of small size, the complexity is polynomial. 

The Topological Environment technique allows the analysis of networks in biochemical modules, 
or their geometric equivalents in local toric charts or varieties. 
In particular, the complexity remains polynomial whenever the neighboring molecules of an 
$X_{i}$ S-system equation involve only a few molecules. 
If the number of neighbors is large, it is always possible to divide the analysis into smaller toric varieties. 
The topological technique then makes it possible to glue these varieties into a global variety, 
while combining the Hilbert bases through a standard matrix sum. 
For more complex geometries, the Minkowski sum is applied 
(see \cite{BSturmfels1996}, \cite{GEwald1993}), thereby reducing the overall complexity of computing Hilbert bases.


\section{Conclusions}

We compare the plots of \textbf{quantitative phenotypes} obtained using the Topological Environment approach with those presented in the analysis by Savageau \cite{LomnitzSavageau2014}. Although differences in concentration magnitude are observed in the \textbf{ logarithmic space}, the behavior of the variable \( X_{4} \) consistently exhibits \textbf{oscillatory dynamics} across different transcriptional control modes.\\

It is important to note that both sets of simulations rely on specific choices of \textbf{free kinetic parameters} and \textbf{initial conditions} for the underlying system of ordinary differential equations. However, the Topological Environment framework provides an alternative and complementary approach to extract equivalent regulatory insights from the original model and offers a pathway to refine the mathematical representation of the system. \\

\textbf{It is also important to note that the solution for the dynamics of the regulator
$X_{4}$ addresses only a subset of S-system equations, without including the differential analysis of the variables $X_{1}$, $X_{A}$, and $X_{2}$, which represent proteins and mRNA transcripts. This distinction arises from a key feature of local analysis enabled by the topological environment methodology. In summary, the local symmetries and properties captured by the Hilbert basis allow us to resolve complex dynamics in smaller geometric pieces, as mentioned earlier, while simultaneously dividing global dynamics into local dynamics.}\\


We have demonstrated that, by employing an alternative approach grounded in \textbf{toric algebraic geometry}, it is possible to obtain results comparable to those derived from the classical analysis of gene circuits as modeled by Lomnitz and Savageau. This is achieved by focusing solely on the \textbf{exponent-space} associated with the underlying mathematical expressions.\\

Within this framework, we uncover \textbf{a geometric perspective} of genetic-metabolic networks, most notably the \textbf{ fixed points on the algebraic torus}. Interestingly, these invariants are \textbf{ independent of the kinetic orders and parameters}, which is a highly relevant outcome. It enables the modeling of genetic and metabolic networks even in the absence of detailed kinetic information, thereby opening new possibilities for the analysis of \textbf{large-scale gene networks}.\\

Furthermore, this approach provides a principled method for identifying \textbf{dominant fluxes}, a fundamental challenge in the study of such networks, especially when dealing with high-dimensional nonlinear dynamics in molecular networks.
%%%%%%%%%%%%%%%%%%%%%%%%%%% DISCUSSION RECOMMENDEN PAPERS %%%%%%%%%%%%%%%%%%
\begin{document}

Once the complete ideas about the methodology in the Topological Environment have been presented, it is relevant to discuss the contributions and challenges introduced by this technique in contrast with some of the models mentioned in the introduction of this work. 

One of the main contributions of the S-system formalism is its relationship with hidden symmetries, which can be resolved through the action of Lie Groups on S-systems. The scaling of variables to map S-systems into homogeneous coordinates for reparametrization was not directly applied in the monomial map used in this work. However, this transformation can be naturally captured, since toric varieties allow embedding non-regular varieties into regular or smooth algebraic varieties. The original space where any S-system has been embedded in toric geometry is the algebraic torus $\mathbb{C}^{*}$. Its coordinates belong to a non-regular toric space given by $\mathbb{C}^{n-r} \times Cone^{r}$, where this space contains an apex or singular space called the non-regular cone $Cone(r)$.  

The Hilbert basis and Hironaka’s theorem guarantee that in a finite number of steps this singular space can be transformed into a regular or smooth toric variety. This process is called a \textbf{Resolution of Singularities} or \textbf{Blow Up}. A practical algorithm to resolve singularities consists of applying a finite number of times the Hilbert basis algorithm through the monomial map and rewriting the equations into the monomial transformation. The resulting space, once the non-regularity is resolved, is a homogeneous space given by $\mathbb{C}^{n-r} \times \mathbb{CP}^{r}$, where the non-regular geometry has been replaced by homogeneous coordinates in the \textbf{complex projective space} $ \mathbb{CP}^{r}$.  

Thus, the scaling of S-system equations can be resolved by this process. It is worth noting that the Hilbert basis is a unimodular transformation that belongs to a Lie Group. In these smooth varieties, local transformations are solved through the Jacobian matrix of $\mathbb{C}^{n-r} \times \mathbb{CP}^{r}$, defined by the Hilbert basis with $ \det(H)\neq 0$. With this fact, one may address the conjecture for models where $ \det(G-H) =0$. It is also important to remark that once the non-regularity of toric varieties has been resolved, there exists a unique fixed point on the algebraic torus, namely $(1,1,\ldots,1) \in \mathbb{C}^{*}$, which recovers a classical result of the analysis in S-systems.\\

For the cases of Statistical Learning Machines or Learning Manifolds (see \cite{rev21}, \cite{rev22}, \cite{rev23}, \cite{rev24}, \cite{rev25}), such as multifractal analysis of stochastic large-scale networks \cite{rev21}, a Fractal Power-Law is defined by  $A_{\alpha}(d,t)=\dfrac{d^{\alpha}_{i}}{\sum^{N(t)}_{i}},$ where $d^{\alpha}_{i}$ is a parameter related to the connectivity degree of hub nodes and $N$ is the number of neighbors varying over time. Here, the likelihood probability is maximized through a stochastic Maximum Expectation Algorithm. In this case, the performance of the statistical machine is improved using the Kullback–Leibler distance $KL(p, q)$, where $p$ is a prior probability distribution modeling the training dataset, and $q$ is the posterior parametric distribution learned by the statistical model $A(d,t)$.  

Most statistical learning models use metrics such as Kullback–Leibler distance, mutual information $I(p,q)$, Fisher information matrix, among others, to improve the learning curve and training parameters. It has been extensively studied by S. Watanabe that such models in their parameter space contain a large number of singularities in high dimensions. He proved that these models are singular in parameter space (see \cite{Watanabe1}, \cite{Watanabe2}, \cite{Watanabe3}, \cite{Watanabe4}). The central issue lies in the fact that the prior distribution $p$ and the posterior distribution $q$ are not bijective in their domain spaces. That is, the set defined by $p(X)\neq q(x| W)$ is not injective, and hence there is no one-to-one correspondence between the parameter spaces. Such statistical learning machines are called non-identifiable.   

The set $\{p(X) = q(x| W)\}$, when the model is identifiable, can be studied as an algebraic set. The resolution of singularities in these high-dimensional spaces can be addressed using the technique of Blow Ups and Hilbert basis \cite{MPCastilloOscar2017} (manuscript in preparation). This approach is further justified by Hironaka’s theorem \cite{MFAtiyah1970}. It is important to mention, that the author has worked new techniques using Hilbert basis generated through of Statistical Machines such as Boltzmann distribution to detect interaction, protein-protein and transcription factor binding to DNA, with good results with this approach, see \cite{Monoids}. \\

Finally, the equilibrium point studied in \cite{rev12}, where a relationship between the stoichiometric space associated with general mass-action balance of a set of reactions and the signed lattice cones in the space of kinetic orders is elucidated, may represent a formal analysis of the genotype–phenotype relationship modeled by genetic and metabolic networks. New properties could be elucidated for the analysis of the Topological Environment and the monomial transformations governed by the Hilbert basis.  

%\end{document}


%%%%%%%%%%%%%%%%%%%%%%%%%%%%%%%%%%%%%%%%%%%%%%%%%%%%%%%%%
\section*{Acknowledgments}
Marco Polo Castillo Villalba is a doctoral student from the Programa de Doctorado en Ciencias Biomédicas, Universidad Nacional Autónoma de México (UNAM) and has received the
CONAHCYT fellowship 754050.
We acknowledge funding from UNAM and from the National Institutes of Health (grant number 5R01GM110597-03). Also, I give huge thanks to Michael A. Savageau, for all the discussions and technical suggestions, during my scholar stay with him, Julio Collado-Vides, Pedro Miramontes and Yair Romero. 
%%%%%%%%%%%%%%%%%%%%%%%%%%%%%%%%%%%%%%%%%%%%%%%%%%%%%%%%%%%%%
\section*{Declaration of Interest Statement}

The author declares that he has no personal, financial, professional, or 
academic conflicts of interest.
%%%%%%%%%%%%%%%%%%%%%%%%%%%%%%%%%%%%%%%%%%%%%%%%%%%%%%
\section*{Declaration of AI-Assisted Writing}
This manuscript benefited from the use of generative AI tools (e.g., OpenAI's ChatGPT) during the writing and editing process. These tools were used to improve the grammar, clarity and structure of the text. All technical content, data analysis, and scientific interpretations were developed and validated solely by the author. The author assumes full responsibility for the integrity, originality, and accuracy of the work.


\input amsart-template_topenv.bbl

%\input amsart_template.bbl
\end{document}


\section{References}


%\subsection{The list of references}

%References should be listed at the end of the main text in the order in which they are first cited in the text. Include all author names with surname first, then initials. Separate multiple author names with semicolons, and multiple editor names with commas. If a reference has more than ten authors, list the first ten names followed by `et al.'. The following list shows some sample references prepared in Taylor \& Francis' Reference Style P.

\begin{thebibliography}{99}

\bibitem{Gou02}%01
Gould, S.J. \emph{The Structure of Evolutionary Theory}; Belknap Press:
  Cambridge, MA, 2002.

\bibitem{LeC03}%02
Le~Couteur, P.; Burreson, J. \emph{Napoleon's Buttons: How 17 Molecules Changed
  History}; Jeremy P. Tarcher/Putnam: New York, 2003; pp 32--47.

\bibitem{Ban75}%03
Bandy, A.R., Ed. \emph{The Chemistry of the Atmosphere: Oxidants and Oxidation
  in the Earth's Atmosphere}; Royal Society of Chemistry: Cambridge, UK, 1995.

\bibitem{Sah04}%04
Saha, B.C., Hayashi, K., Eds. \emph{Lignocellulose Biodegradation}; ACS
  Symposium Series 889; American Chemical Society: Washington, DC, 2004.

\bibitem{Hil90}%05
Hillman, L.W. In \emph{Dye Laser Principles with Applications}; Duarte, F.J.,
  Hillman, L.W., Eds.; Academic: New York, 1990; Chapter~2.

\bibitem{Pul04}%06
Puls, J.; Saake, B. Industrially Isolated Hemicelluloses. In
  \emph{Hemicelluloses: Science and Technology}; Gatenholm, P., Tenkanen, M.,
  Eds.; ACS Symposium Series 864; American Chemical Society: Washington, DC,
  2004; pp 24--37.

\bibitem{Kli04}%07
Kline, R.B. \emph{Principles and Practice of Structural Equation Modeling}, 2nd
  ed.; Guilford Press: New York, 2004.

\bibitem{Leo05}%08
Leone, S.R., McDermott, A.E., Paul, A., Eds. \emph{Annual Review of Physical
  Chemistry}; Annual Reviews: Palo Alto, CA, 2005; Vol.~56.

\bibitem{Wib74}%09
Wiberg, K. In \emph{Investigations of Rates and Mechanisms of Reactions};
  Lewis, E.S., Ed.; Techniques of Chemistry, Vol.~VI, Part~I; Wiley: New York,
  1974; p 764.

\bibitem{McG02}%10
\emph{McGraw-Hill Encyclopedia of Science and Technology}, 9th ed.;
  McGraw-Hill: New York, 2002; 20 vols.


\section{Appendices}


\appendix


\subsection{Via e-mail}

This article template, the \texttt{interact} class file and the associated open-source \LaTeX\ packages are also available via e-mail. Requests should be addressed to \texttt{}, clearly stating for which journal you require the template and class file.

\end{document}
