## Supplementary figures and images for "Topological Environment in Genetic and Metabolic Networks"

### Circuit1dpi.png

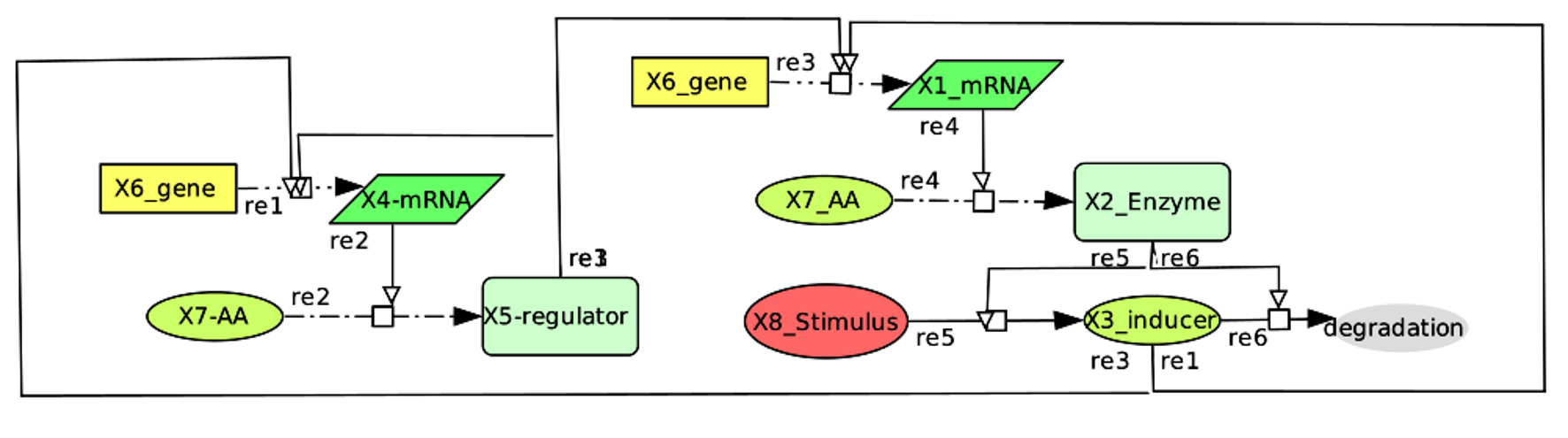

### Circuit2dpi.png

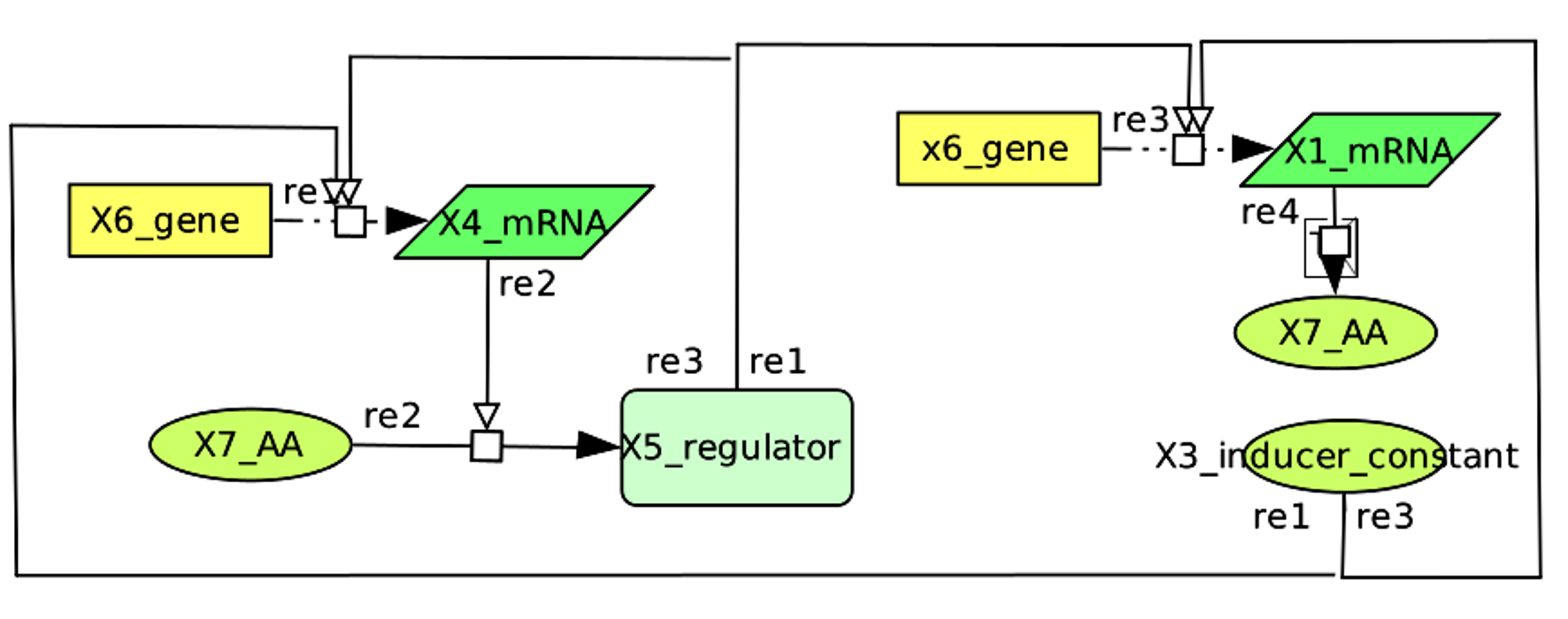

### Circuit3dpi.png

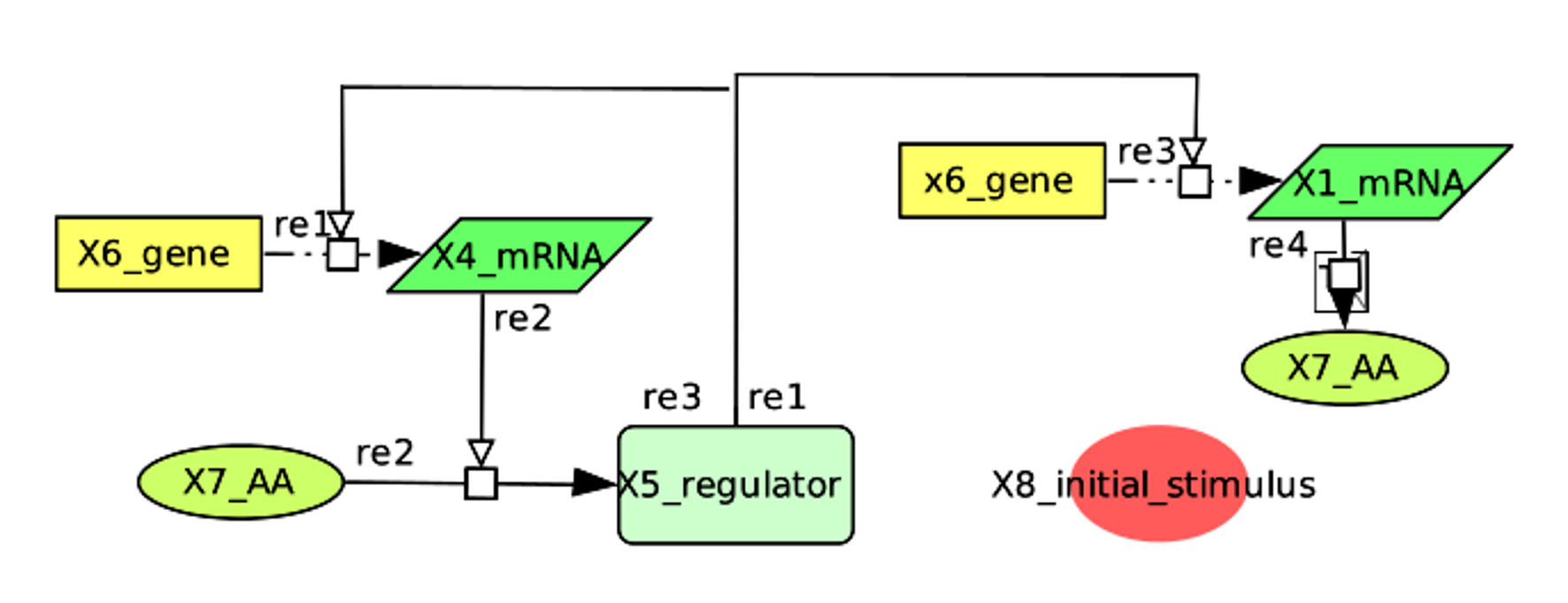

### Circuit4dpi.png

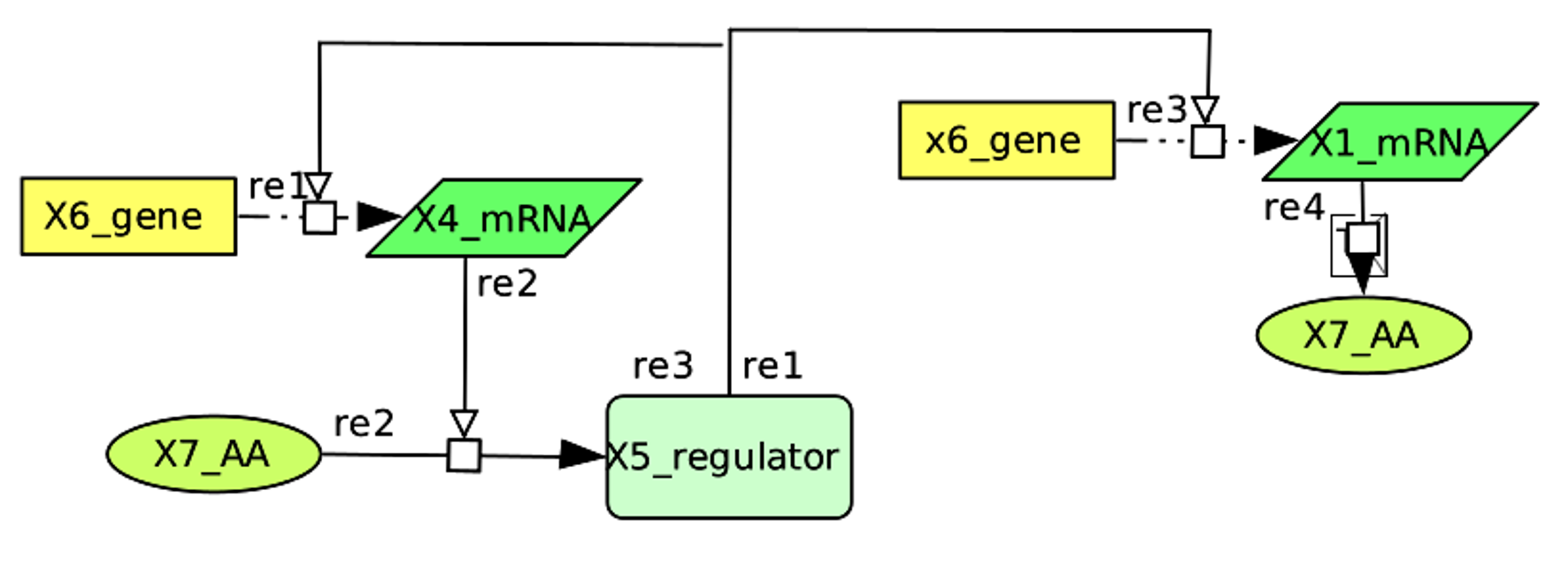

### Circuit5dpi.png

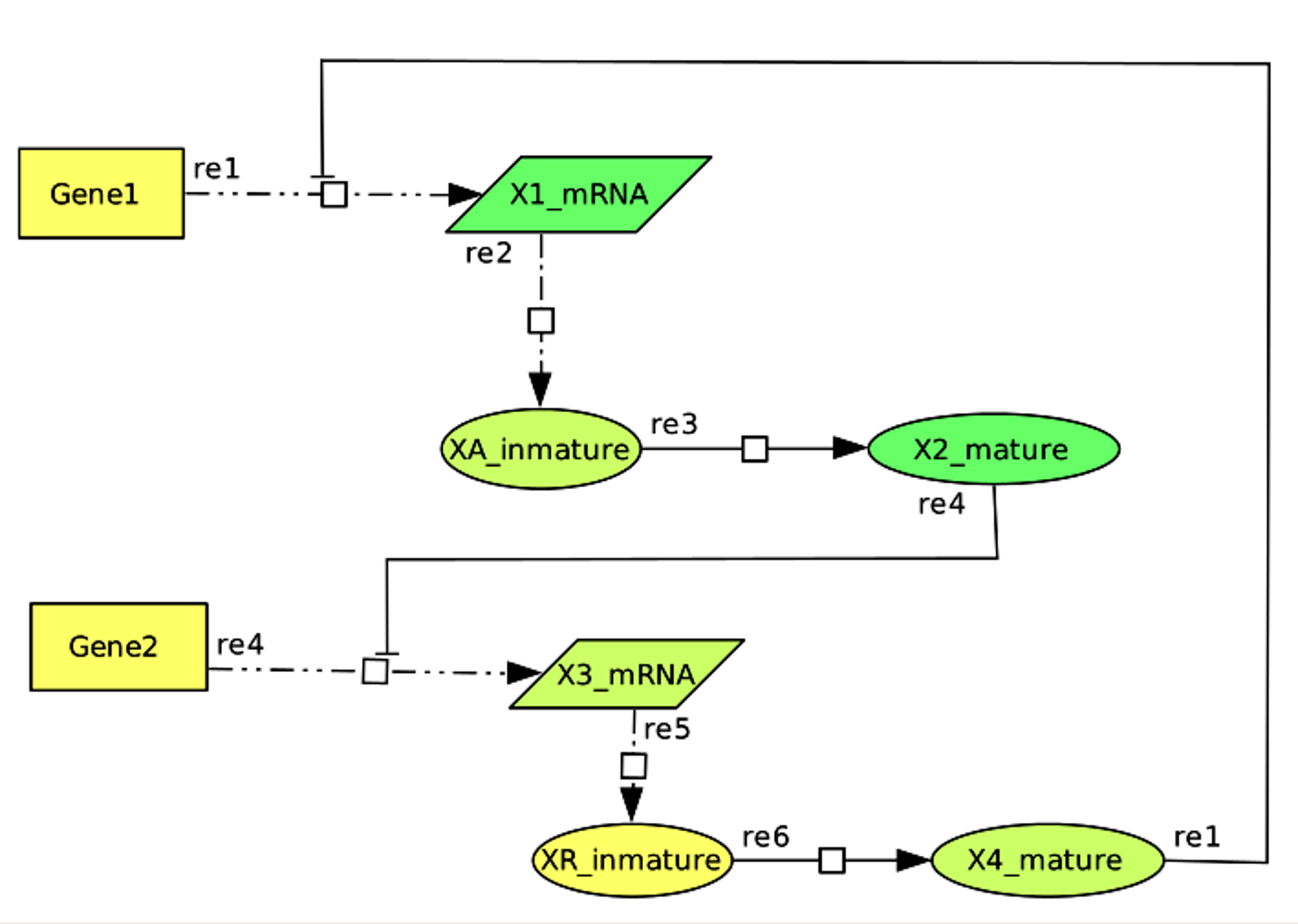

### Circuit6dpi.png

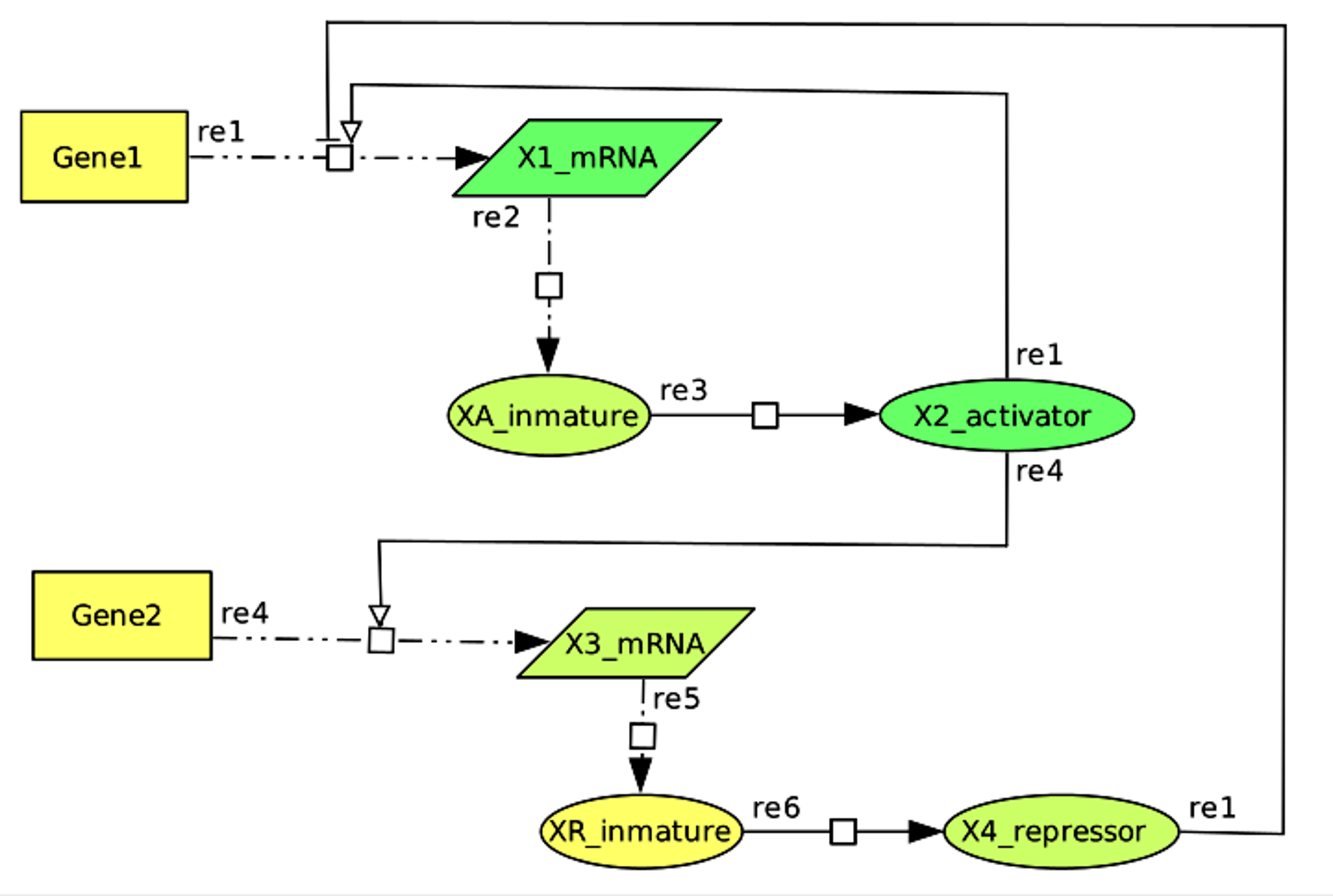

### Circuit7dpi.png

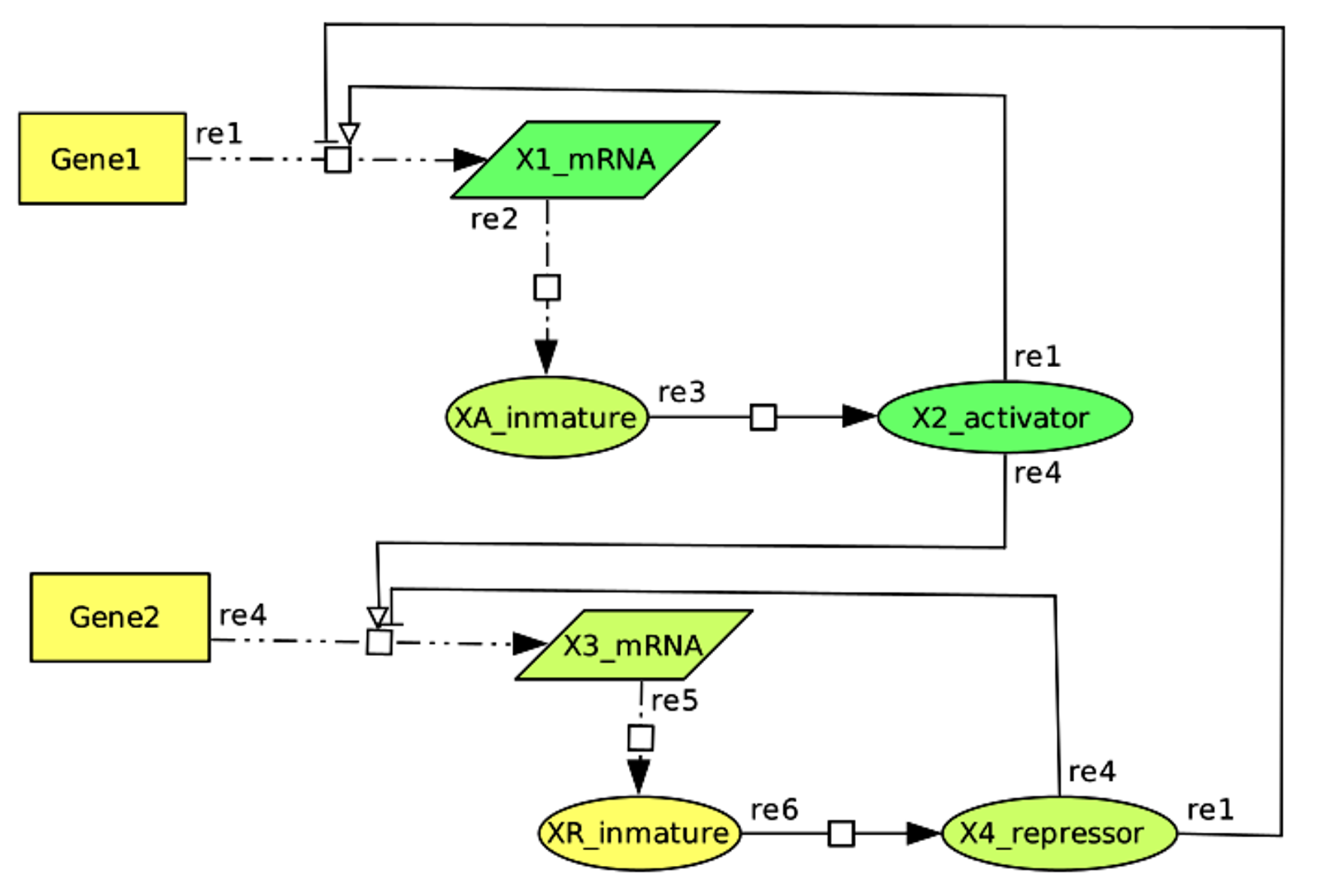

### Circuit8dpi.png

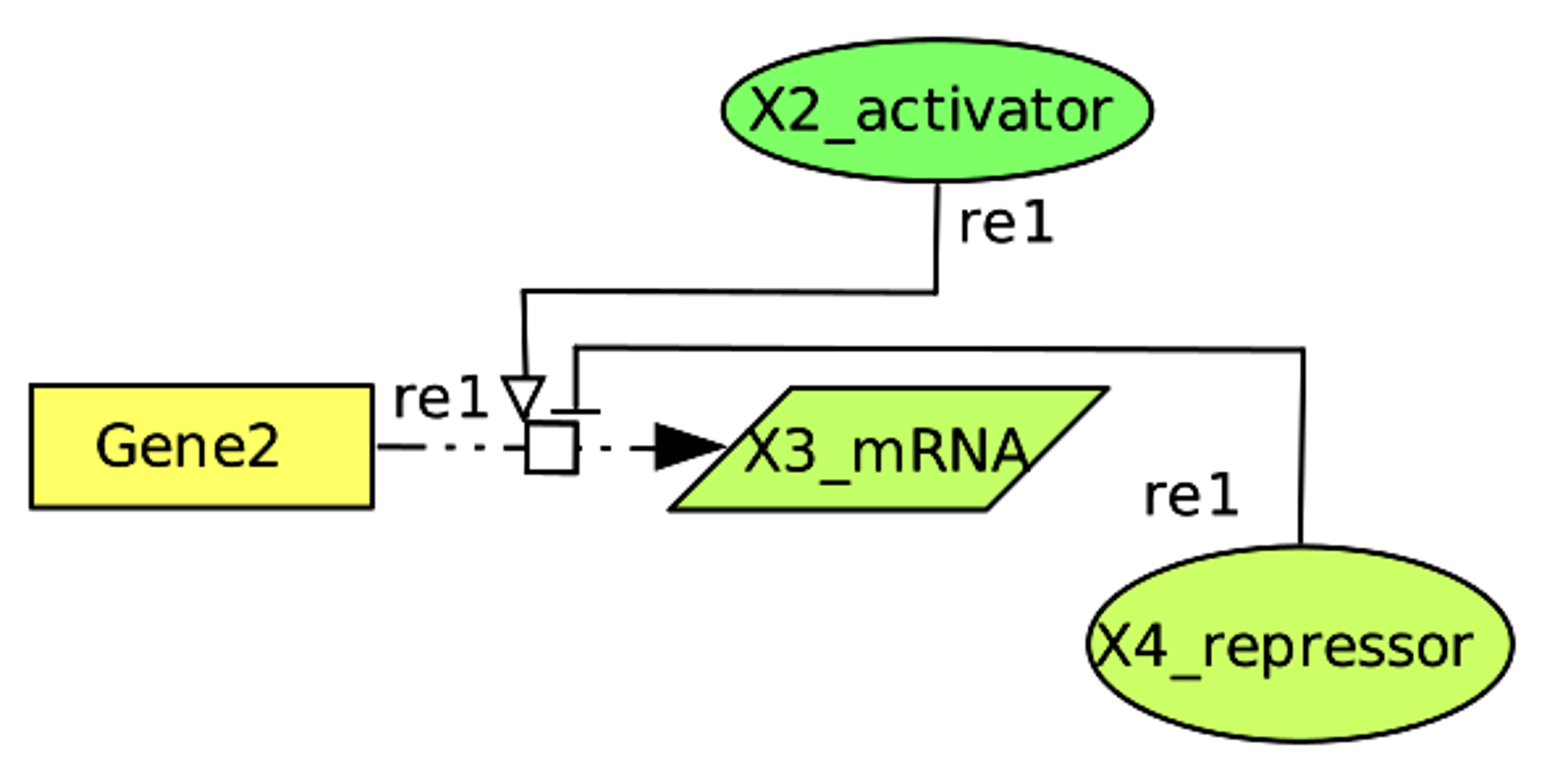

### Circuit9dpi.png

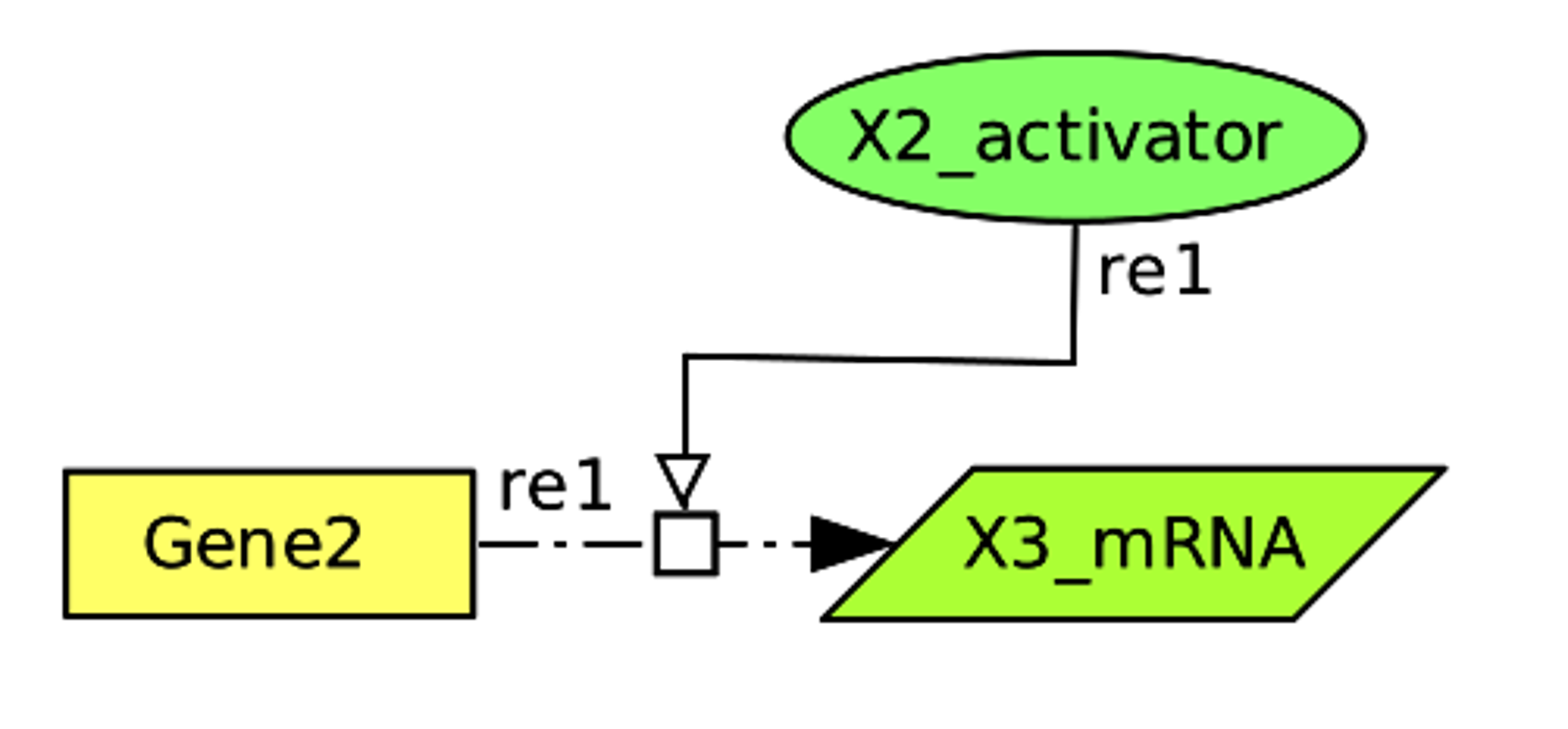

### Circuit10dpi.png

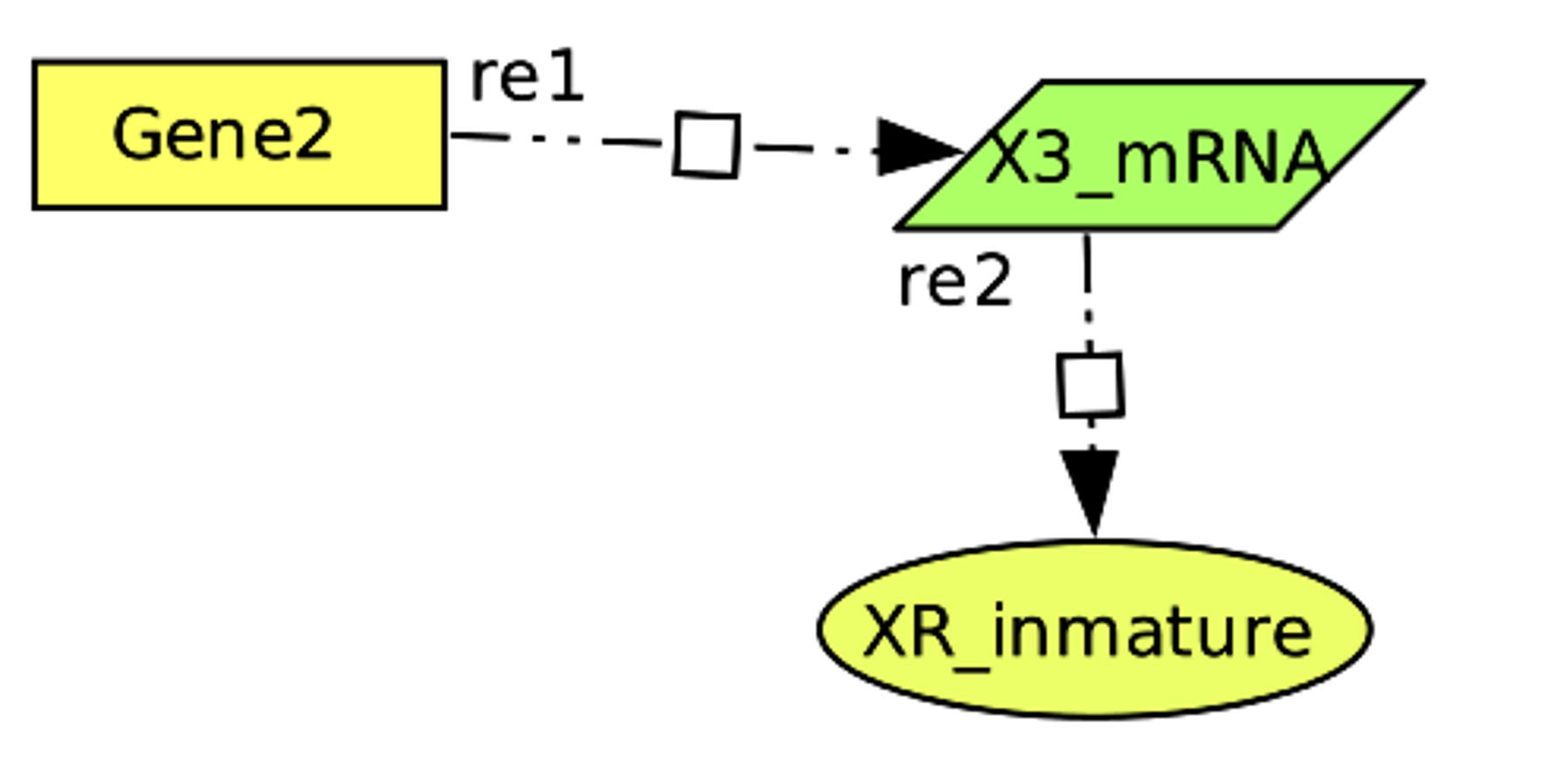

### Simulations.png

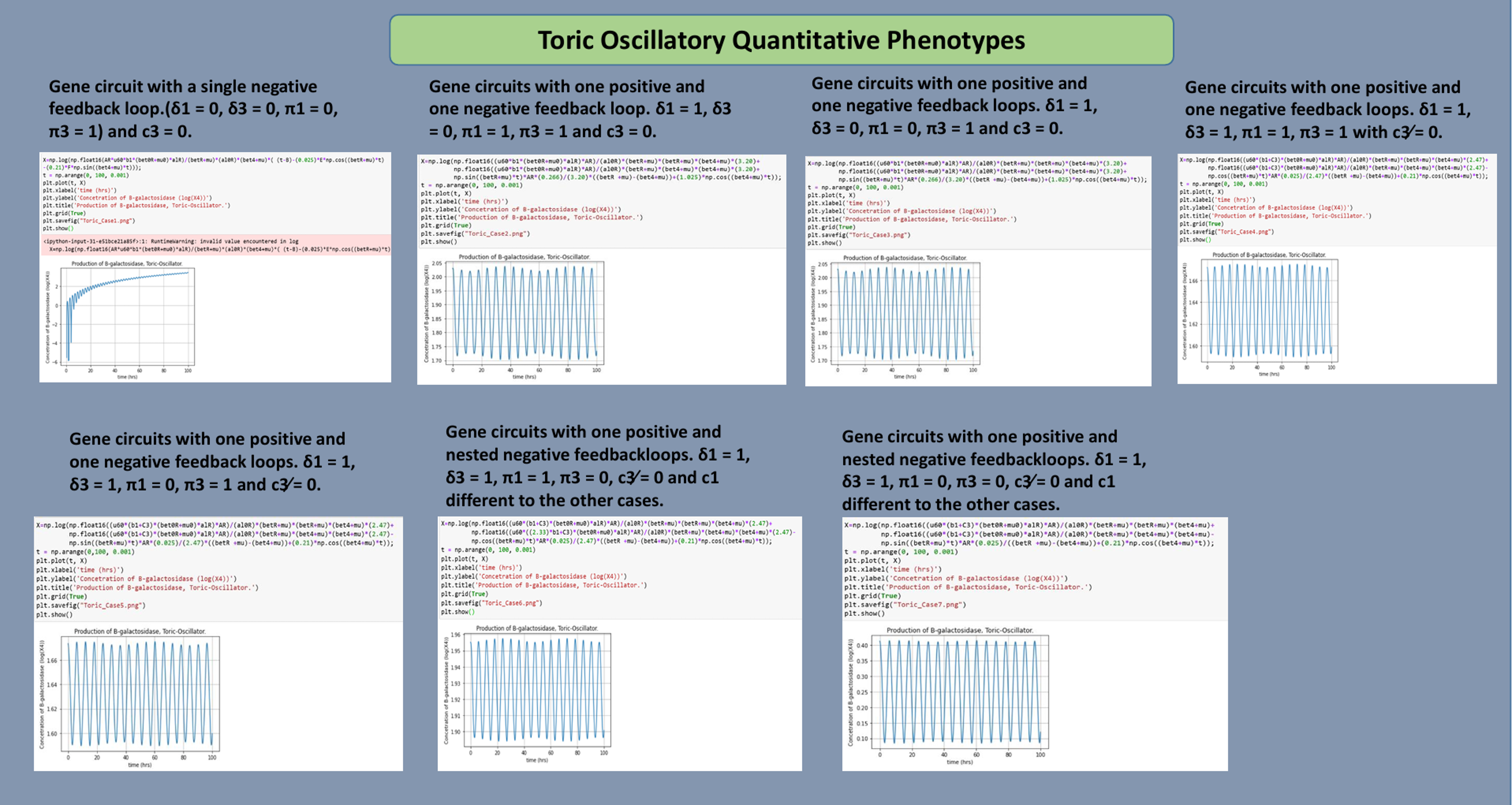

### top_env1dpi.png

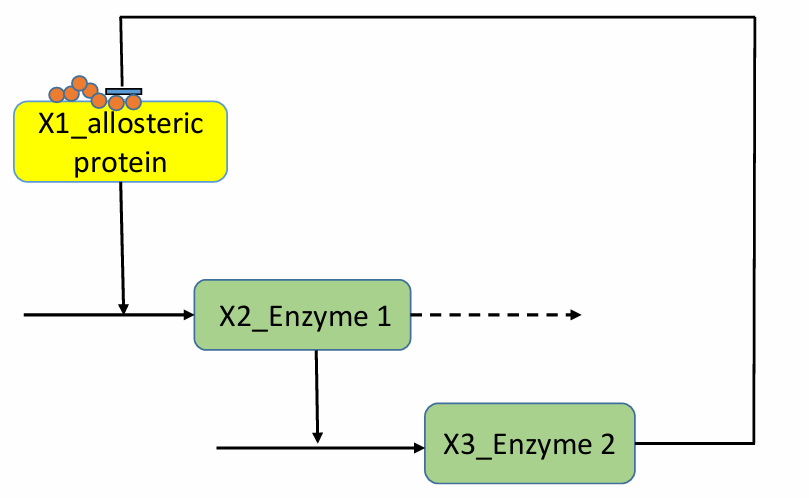

### top_env5dpi.png

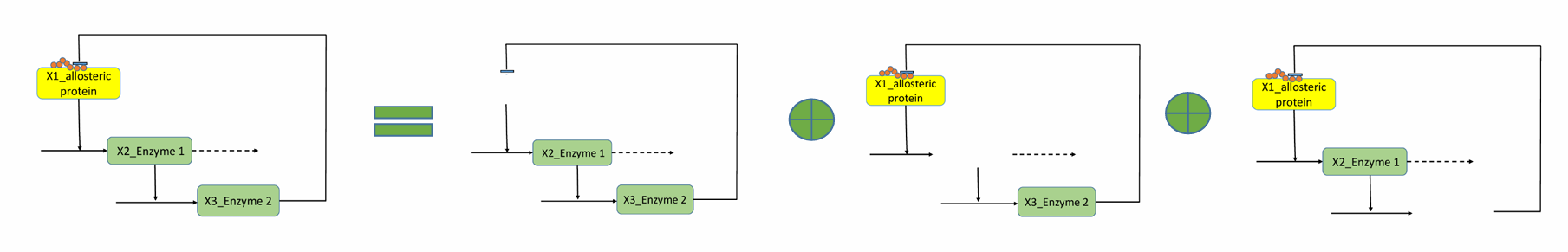
